# Breast cancer ovarian metastases show increased activity of GPCR pathways

**DOI:** 10.64898/2026.07.30.741805

**Authors:** Laura Savariau, Ye Qin, Osama Shah, Ahmed M Basudan, Chris Merkel, Saumya D. Sisoudiya, Smruthy Sivakumar, Ethan S. Sokol, Olivia McGinn, Zheqi Li, Tiantong Liu, Nilgun Tasdemir, Pooja Tallapaneni, Lan Coffman, Esther Elishaev, Jennifer M. Atkinson, Peter C. Lucas, Adrian V. Lee, Steffi Oesterreich

## Abstract

Treatment resistance and metastases occur in 10–20% of patients with invasive lobular carcinoma (ILC), the most common special histological subtype of breast cancer. ILC metastasizes to the ovary more frequently than no special type (NST) tumors, also known as invasive ductal carcinoma (IDC). To characterize the genomic landscape of breast cancer ovarian metastases, we analyzed 15,613 local breast cancers, 22,010 non-ovarian metastases, and 246 ovarian metastases sequenced using FoundationOne®CDx or FoundationOne® assays. Ovarian metastases had enriched *CDH1*, *PIK3CA*, and *TBX3* mutations and depleted *TP53* and *MYC* alterations relative to local breast cancers, with additional depletion of *ESR1* mutations compared to non-ovarian metastases. *CDH1* mutations were less frequent in ovarian metastases (47%) than local ILC (81.3%), with reduced 16q loss (64% vs 84%), indicating that ovarian metastases also arise from non-ILC tumors. We extended these findings to a UPMC cohort of 27 ovarian metastases (13 ILC, 8 IDC, 6 mixed ductal-lobular carcinoma) with patient-matched primary tumors in most cases. In both cohorts, patients with ovarian metastases were significantly younger than those with other metastatic sites. In the UPMC cohort, the most frequent mutations were in *PIK3CA*, *CDH1*, *KMT2C*, *FOXA1*, and *RUNX1*. Transcriptomic analysis identified upregulated G protein-coupled receptor (GPCR) pathways, including metabotropic glutamate receptor signaling. Functional studies showed that calcium-sensing receptor (CaSR), a GPCR overexpressed in ovarian metastases, drives MEK/ERK-dependent migration and F-actin reorganization in ILC cell lines, enhanced by estrogen and blocked by calcilytic, MEK, or anti- estrogen treatment. Our findings inform future therapeutic targeting of ovarian metastasis.

## Introduction

In addition to the “no special type” (NST) breast cancer, also called invasive ductal carcinoma (IDC), there are special types, among which invasive lobular carcinoma (ILC) is the most common. ILC represents 10-15% of all invasive breast cancer [1, 2], and comprises 28,000-42,000 new annual breast cancer diagnoses in the United States [3]. The hallmark of ILC is the genomic loss of E-cadherin *(CDH1)* which leads to dis-cohesive and single cell file growth pattern and renders ILC difficult to be detected via imaging [4]. Compared to NST/IDC, ILC tumors are larger at presentation and often multifocal. 94% of ILC are estrogen receptor positive (ER+), 83% are of the luminal A molecular subtype of breast cancer, and they typically display a low proliferation rate [5, 6]. Despite these favorable prognostic factors, patients with ER+ ILC do not have a better outcome compared to ER+ IDC, and in contrast the rates of late recurrences are higher [5, 7, 8].

In 1984, Harris *et al* reported that ILC metastasizes more frequently to the gastrointestinal tract, peritoneum, and urogenital organs including the ovaries compared to IDC [9]. Two decades later, Arpino *et al* undertook a comprehensive review of the metastatic spread and clinical outcome of 4,140 ILC and 45,169 IDC cases with a median follow-up period of 87 months [10]. In their analysis, ILC was three times more likely to spread to the ovaries, peritoneum, and gastrointestinal (GI) system compared to IDC (6.7% vs 1.8% in IDC), whereas lung, distant lymph nodes, and CNS metastases were observed more frequently in IDC cases. We have previously reported on ophthalmic metastases which is enriched in patients with ILC [11]. Subsequent studies have confirmed these observations on differences in metastases between IDC and ILC at varying frequencies [8, 11–13]. Of note, there is a general underreporting of metastases to unique sites such as the ovary as such data is frequently not captured in registries.

Metastatic breast cancer comprises 3-38% of all ovarian tumors [14–17]. Aside from the association with lobular histology, ovarian metastasis from the breast has been reported to be associated with large tumor size, lymph node involvement, and high histologic grade [16]. Ovarian metastases also occur more frequently in younger, pre-menopausal patients [17]. The median overall survival of patients with ovarian metastases from breast cancer was reported to range from 16-38 months, and the 5-year survival rate is 6-26% (reviewed in [17]). Ovarian metastases from ILC are frequently misdiagnosed as primary ovarian cancer on imaging, which can lead to unnecessary gynecologic surgery. Percutaneous biopsy of an accessible metastatic site has been shown to correct the diagnosis and direct patients toward systemic rather than surgical management[18].

Cytoreductive surgery is the primary management strategy, although it is unclear which patient populations benefit the most from these procedures [14, 17]. Targeted therapies to specifically treat breast cancer ovarian metastases are currently unavailable and are urgently needed.

Although E-cadherin loss is thought to be a driver of cancer metastasis, its role in ILC metastasis remains to be investigated [19–21]. In general, there is a great need to better understand the biology regulating the dissemination of lobular tumor cells, especially considering the unique organ tropism. Here, we present the genomic landscape of 246 ovarian metastases sequenced through FoundationOne (F1Cdx and F1) assays. In addition, we report characterization of clinicopathological features and multiomics analysis of a UPMC cohort of 12 pairs of primary breast tumor and ovarian metastasis as well as 14 orphan breast cancer ovarian metastases. We identified a series of glutamate receptors (GluR) including metabotropic GluR G-protein coupled receptor (GPCR) as being up-regulated in ovarian metastases. Another GPCR, the Calcium Sensor Receptor (CaSR) [22] previously associated with bone metastasis [23–25] was also upregulated. Our functional studies show that CasR mediates migration of ILC cells in an ER and MEK/ERK-dependent fashion. In addition, we uncovered multiple pathways which should be further explored for potential therapeutic targeting of breast cancer ovarian metastasis.

## Results

### Mutational and chromosomal landscape of breast cancer ovarian metastases in a large clinical sequencing cohort

To characterize the genomic landscape of breast cancer metastasis to the ovary, we analyzed all classes of alterations including mutations, rearrangements, and copy number changes from a large dataset comprising 15,613 local breast cancer tissues (of which 1,350 were from patients with ILC), 22,010 non-ovarian metastases, and 246 ovarian metastases, all of which underwent tumor-only sequencing using the (F1Cdx) or (F1) assays.

Patients with ovarian metastases were younger than those in the comparator groups, with a median age of 52 years, compared to 60 years for non-ovarian metastases, 57 years for local breast cancer of any subtype, and 62 years for local ILC (*P* < 0.0001; **Table 1**).

**Table 1.** Overview of FMI Cohort. Median age of patients diagnoses with OvM was younger than non-ovarian metastases. There were fewer HRDsig+ tumors in ILC breast samples and ovarian metastases compared to all local breast samples and non-ovarian metastases, respectively.

|  | Local breast samples | Local ILC samples | Ovarian samples | Non-ovarian samples | p-value |
| --- | --- | --- | --- | --- | --- |
| # of samples | 15,613 | 1,350 | 246 | 22,010 |  |
| Median age, y (IQR) | 57 (47-67) | 62 (53-70) | 52 (45-60) | 60 (51-68) | $P < 0.0001$ |
| %HRDsig +ve (# +ve/#assessable) | 17.6% (2667/15114) | 6.6% (85/1288) | 8.1% (19/234) | 14.3% (3026/21223) | $P < 0.0001$ |
| Median TMB (IQR) | 2.5 (1.3-4.3) | 2.5 (1.3-3.8) | 2.5 (1.3-4.8) | 2.6 (1.25-5) | $P < 0.0001$ |
| MSI status, n (%) | | | | | $P < 0.001$ |
| MSS | 14,812 (95%) | 1,311 (97%) | 237 (96%) | 21,122 (96%) |  |
| MSI-High | 78 (0.5%) | 6 (0.4%) | 2 (0.8%) | 101 (0.4%) |  |
| MSI-L | 662 (4%) | 32 (2.4%) | 7 (2.8%) | 697 (3.2%) |  |
| Unknown | 61 (0.5%) | 1 (0.06%) | 0 | 90 (0.4%) |  |

Median tumor mutational burden (TMB) was comparable across all four cohorts (range 2.5 - 2.6 mut/Mb), and the proportion of microsatellite stable (MSS) cases was uniformly high (≥95%) in each group (**Table 1**). Homologous Recombination Deficiency Signature (HRDsig) positive cases were less frequent in ovarian metastases (8.1%) compared to non-ovarian metastases (14.3%), and similarly less frequent in local ILC (6.6%) compared to local breast cancer overall (17.6%) (*P* < 0.0001; **Table 1**).

Comparing the prevalence of gene alterations in ovarian metastases to local breast biopsies, we identified enrichment of *CDH1*, *PIK3CA*, and *TBX3* mutations and attenuation of *TP53* and *MYC* gene alterations in the ovarian metastases (**Figure 1A**). A similar pattern emerged when comparing ovarian to non-ovarian metastases, but additional depletion of *ESR1* mutations in ovarian metastases (**Figure 1B**).

**Figure 1.**
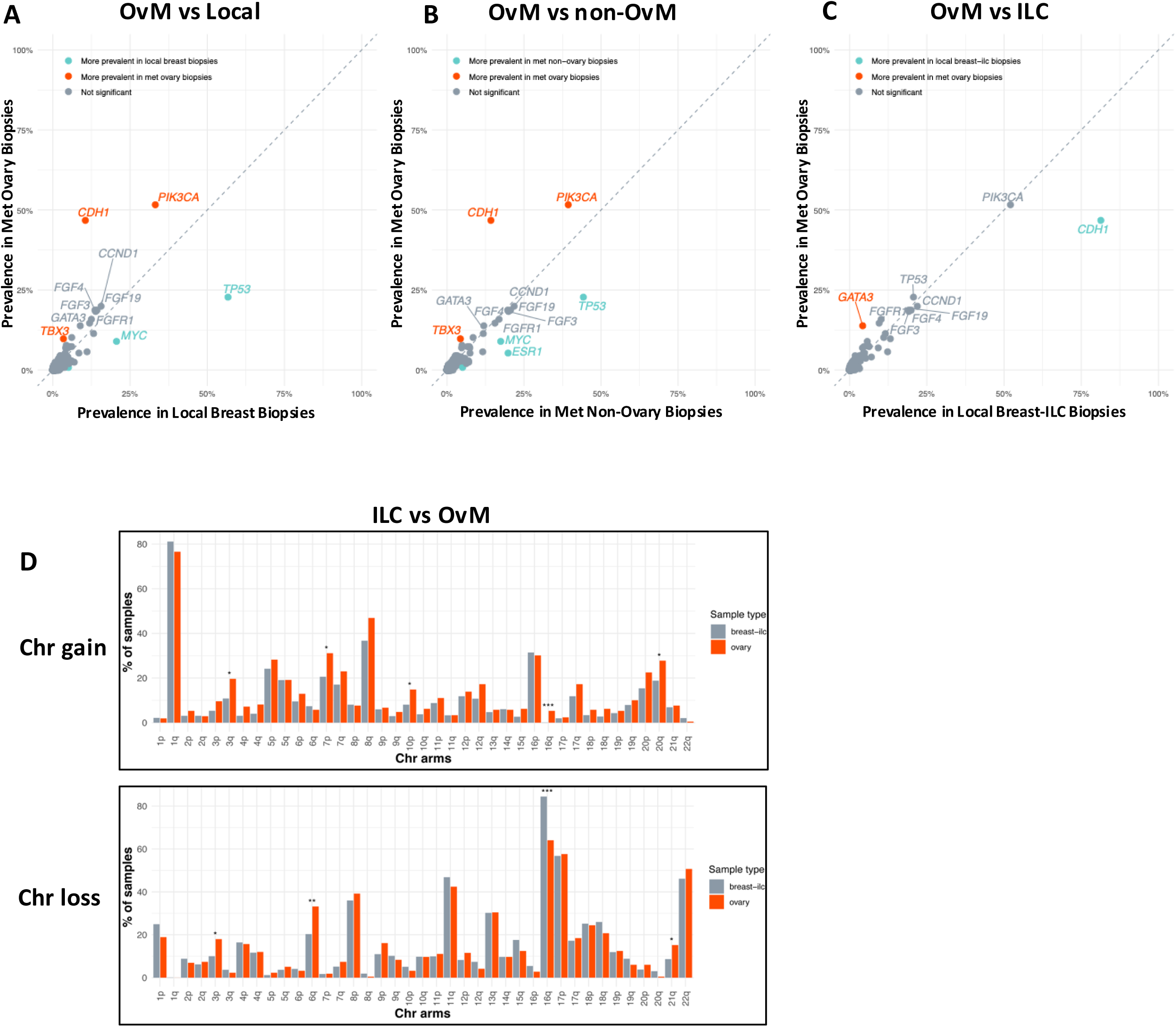
Mutational and Chromosomal Alterations in Ovarian Metastasis. (A–C) Gene-level mutation prevalence in ovarian metastases versus three FMI comparator groups. Each point is a gene, plotting prevalence (percent of samples altered) in ovarian metastatic biopsies (y-axis) against the comparator (x-axis). Genes enriched in ovarian metastases are red, genes enriched in the comparator are cyan, and non-significant genes are grey (A, B) or yellow (C). (A) Versus local breast biopsies (n=15,613). CDH1, PIK3CA, and TBX3 were enriched in ovarian metastases; TP53 and MYC in local breast. (B) Versus non-ovarian metastases (n=22,010). CDH1, PIK3CA, and TBX3 were enriched in ovarian metastases; TP53, MYC, and ESR1 in non-ovarian metastases. (C) Versus local breast ILC biopsies (n=1,350). CDH1 was enriched in local breast ILC; other labeled genes, including PIK3CA, did not differ. (D) Chromosomal arm-level gains (top) and losses (bottom) in local breast ILC (grey) versus ovarian metastases (orange), as percent of samples altered per arm. Asterisks denote significant differences between groups.

*CDH1* mutations were detected in 115 of 246 (47%) ovarian metastases. While this represents a marked enrichment relative to local breast cancers overall (**Figure 1A**), this frequency was lower than the 81.3% reported in local breast ILC (**Figure 1C**), indicating that although *CDH1*-mutant tumors are enriched among ovarian metastases, they can also develop in non-ILC tumors. Consistent with this observation the frequency of arm-level loss at 16q, the chromosome arm harboring *CDH1,* was reduced in ovarian metastases relative to local ILC (64% vs 84%, *P* = 3.4×10⁻⁹) (**Figure 1D**).

### Clinicopathological overview, mutational profiling, and molecular subtyping of UPMC ovarian metastasis cohort

Our single institution (UPMC), retrospective ovarian metastases cohort is comprised of 27 metastatic samples (**Figure 2A**) including 13 paired to 12 primary breast tumor samples and 14 orphan metastatic samples. Using H&E and E-cadherin (and p120) IHC staining, we were able to distinguish between ILC, mixed ductal and lobular carcinoma (mDLC) and IDC subtypes (**Figure 2B**, and deposition in figshare [26]). In total there were 13 ILC, 6 mDLC and 8 IDC ovarian metastasis.

**Figure 2.**
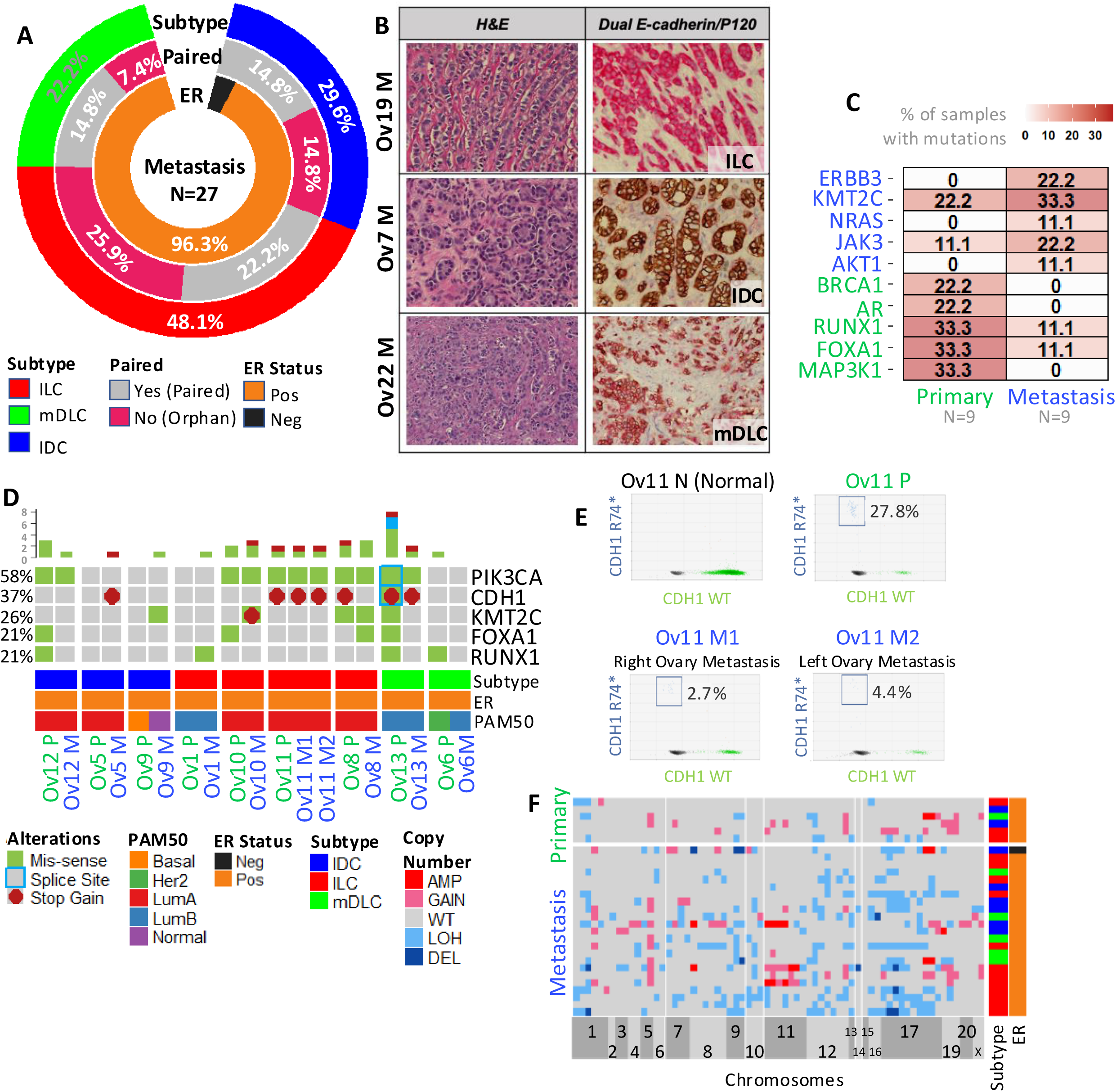
Clinical and pathological features of UPMC ovarian metastasis cohort. (A) Overview of tumor subtype, ER status, and paired status for 27 ovarian metastatic samples. (B) H&E and dual E-cadherin (brown)/P120(red) staining on a representative ovary metastasis from each histological subtype present in our cohort. Images at 20X. M = Metastasis. (C) Genes with largest mutational frequency change between paired primary and metastatic samples. Each number in the table represents relative mutation frequency as “percentages”. (D) Oncoprint showing top 5 most mutated genes in paired primary and ovary metastatic samples. Samples are split by tumor subtype i.e., IDC, ILC and mDLC and tissue type i.e., primary and metastasis. Each column entry is a sample annotated with Patient IDs and P (primary) or M (Metastasis) characters. (E) ddPCR validation of *CDH1* R74* mutation identified in Ov11 P (primary) and Ov11 M1 and M2 (ovary metastasis samples). F) Heatmap showing copy number profiles across primary tumor and ovarian metastasis samples.

In order to assess whether metastasis to ovary was enriched for patients with ILC compared to IDC, we combined our ovarian metastasis dataset with our previously published UPMC breastMETs datasets [27–30] that included information on additional metastatic sites including brain, bone and GI (N=48). Using clinical annotations from the combined dataset, Pearson’s chi square test showed significantly positive association between metastasis to ovary and both ILC and mixed ductal lobular carcinoma (mDLC) subtypes compared to other metastatic sites (**Supplementary Figure S1**).

Analysis of the patient cohort with ovarian metastases revealed a median age at diagnosis of the primary tumor of 43 years, with 78% of patients being premenopausal (**Table 2**). ER positivity was seen in 89% (based on IHC) and 96.3% (based on *ESR1* mRNA expression) of the ovarian metastatic samples (**Figure 2A**, and **Table 2**), and 68% of the samples expressed PR by IHC (**Table 2**). The median disease-free survival (DFS) was 49.5 months. Additional clinicopathological details are provided in **Supplementary Table S1**.

**Table 2.**
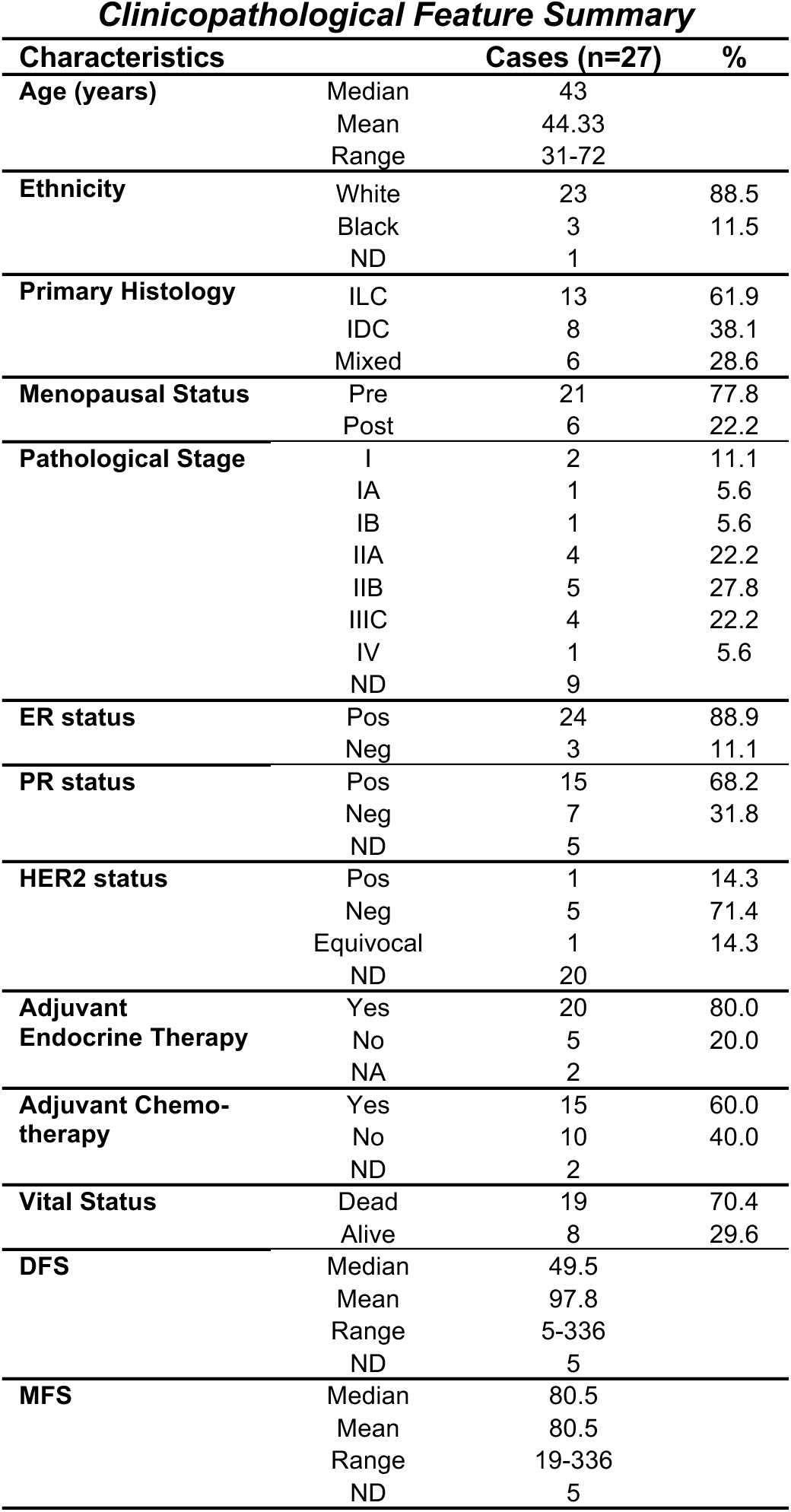
Clinicopathological features of cases included in the study (n=27). ER, PR and HER2 status is based on information from the metastatic samples. Additional and more detailed information the tumors and treatment can be found in **Supplementary Table S1**. “ND” for no data/unknown.

To identify mutations in the ovarian metastases and matched primary tumors, we performed targeted DNA sequencing using the MammaSeq panel previously reported by us [31]. Sufficient DNA for panel sequencing was obtained from 45/49 samples, and all profiled samples passed quality control assessment (average coverage > 100X) (**Supplementary Figure S2A & B**). SNV count before and after filtering is shown in **Supplementary Figure S2C**. The mutations were annotated using cravat and filtered to enrich for somatic variants (**Supplementary Data 1**). Genes with the greatest increase or decrease in number of mutations between intra-patient paired primary and metastatic samples were identified (**Figure 2C**) (comparison between primary tumors and all metastatic samples is shown in **Supplementary Figure S3A**). Compared to primary tumors, paired ovarian metastases had higher number of mutations in *ERBB3, KMT2C, NRAS, JAK3* and *AKT1* and lower number of mutations in *MAP3K1, FOXA1, RUNX1, AR* and *BRCA1,* indicated in **Figure 2C** as percentage of samples mutated. Of the genes covered by MammaSeq, the most frequently mutated genes in the paired primary and ovarian metastasis were *PIK3CA, CDH1*, *KMT2C*, *FOXA1* and *RUNX1* (**Figure 2D**). The top 20 mutated genes across all samples are shown in **Supplementary Figure S3B**. In 2 out of 3 paired samples (Ov11 and Ov13), *CDH1* mutations detected in the primary tumor remained in the ovarian metastases (**Figure 2D**) (*CDH1* gene coverage in MammaSeq panel is shown in **Supplementary Figure S4A**). For example, Ov11 presented with *CDH1* R74* mutation (**Figure 2D**) which was confirmed through inspection of DNA and RNA sequencing reads using the Integrated Genome Viewer (IGV) (**Supplementary Figure S4B**) and by digital droplet PCR (**Figure 2E**). Clinically actionable alterations were identified using OncoKB [32] and PMKB [33] databases. These included mutations in *PIK3CA*, *ERBB2*, *ESR1* and *JAK3* genes (**Supplementary Figure S3C**, and additional details in **Supplementary Data 1**). OncoKB level 3 oncogenic alterations across the samples is shown in **Supplementary Figure S3D**.

We next investigated copy number (CN) profiles of the ovarian metastases using our previously reported dataset generated with targeted gene detection [34]. Overall, CN aberrations were enriched in the metastatic samples compared to primary tumors (**Figure 2F**). Deletions in *JUN, FGFR1, FOXA1, CDKN2A/2B, CDH1, LSMD1* and *TP53* genes and amplifications in *FGFR1, CCND1, FGF19, FADD, PAK1, NF1, RPTOR* and *CTTN* genes were seen in one or more metastatic samples but absent in primary tumors.

We used RNA sequencing to assess transcriptomic alterations associated with ovarian metastasis. All samples passed sequencing quality control assessment with exception of Ov3M which had very low mapped reads (< 10 million) and was therefore removed from the downstream analysis (**Supplementary Figure S5A**). Genotype similarity analysis confirmed that all patient-matched samples were indeed from the same patient (**Supplementary Figure S5B**). Sample-sample similarity analysis based on Euclidean distance (**Supplementary Figure S5C**) and principal component analysis (PCA) (**Supplementary Figure S5D**) showed clustering by tissue type i.e., by primary tumor or ovarian metastasis. Further annotation of the PCA plots revealed associations with ESTIMATE purity scores; of note, we did observe strong correlation between ESTIMATE scores and pathological cellularity values (Pearson correlation coefficient = 0.6 and p < 0.0039) and PAM50 subtypes (**Supplementary Figure S5D**).

The ovarian metastases were predominately of Luminal A subtype (**Figure 3A, B**). We examined PAM50 subtype between patient-matched primary and metastatic disease, given previously described subtype switching associated with drug resistance and invasion in a subset of breast tumors [35–38]. PAM50 switching occurred in 4 primary (P) and metastatic (M) pairs: Ov9, Ov5, Ov13 and Ov1 (**Figure 3A**, highlighted with bolded row names). We observed more diverse PAM50 distribution in IDC and mixed ductal lobular metastases compared to ILC metastases (**Figure 3B**), however the numbers are too small to draw solid conclusions.

**Figure 3.**
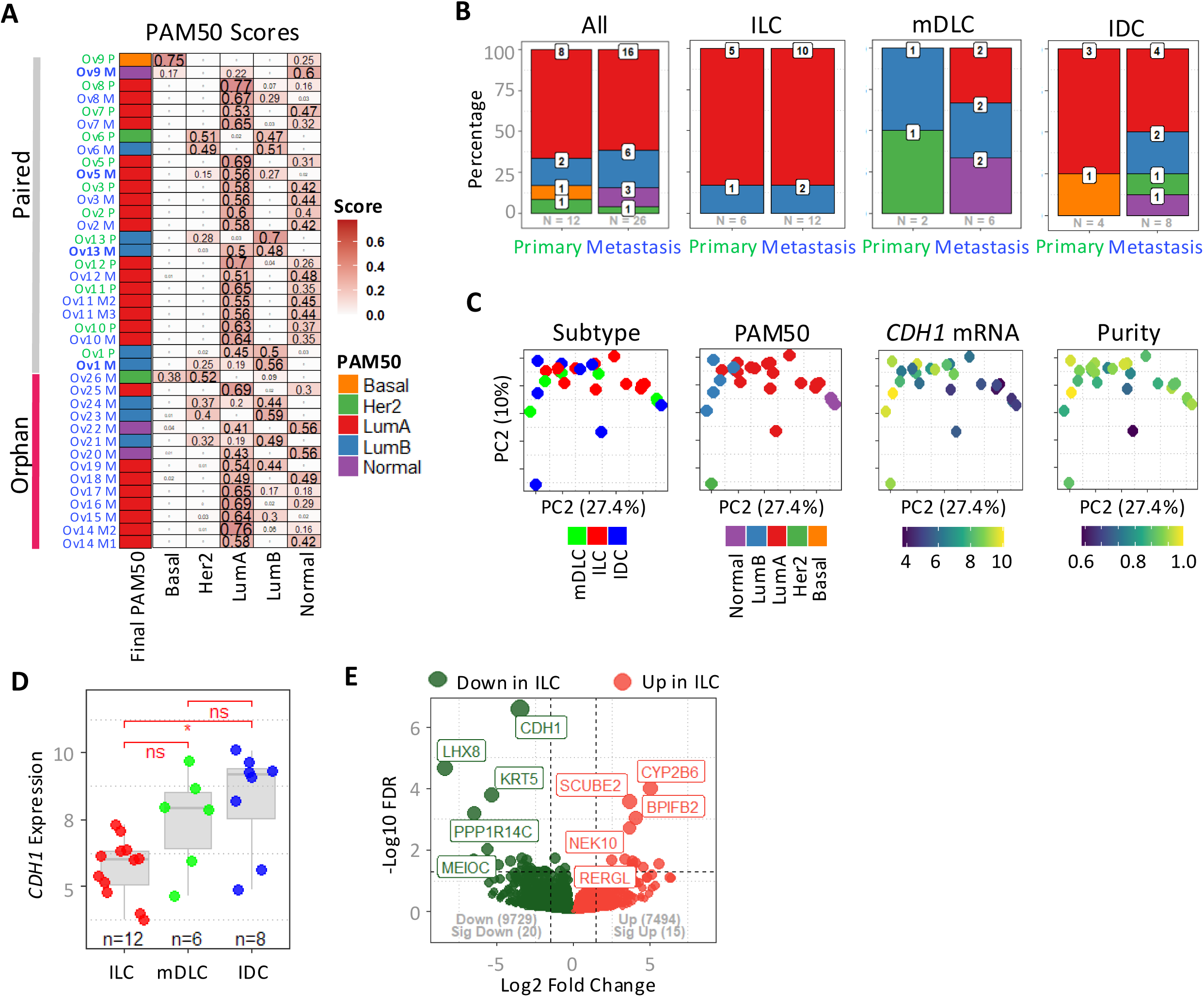
PAM50 subtyping and molecular landscape of ovary metastasis. (A) PAM50 subtype scores across ovary metastasis cohort. Each row entry is a unique sample annotated with Patient IDs and P (primary) or M (Metastasis) characters. Paired and orphan metastatic samples are shown on top and bottom, respectively. Metastatic samples showing PAM50 switching have bolded row entry names. Subtype switching was defined as at least one of the PAM50 subtype call probability scores between paired primary and metastatic samples differed by 0.25 or 25%. “Assignment” column shows the most dominant PAM50 subtype i.e., one with the highest score per sample. Individual PAM50 subtype call probability scores are shown in remaining columns labeled as “Basal”, “Her2”, “LumA”, “LumB” and “Normal”. (B) PAM50 subtype distribution in primary tumor and ovary metastasis in whole cohort and across histological subtypes. (C) PCA analysis of ovary metastasis samples annotated by primary tumor histological subtype, PAM50, *CDH1* expression and Purity. (D) *CDH1* expression differences between ILC, IDC and mDLC ovary metastasis. E) Volcano plot of DEGs in ILC vs IDC ovarian metastatic samples. Statistical comparison in panel D was performed using wilcoxon signed-rank test. “*” denotes p value <= 0.05. Pathway enrichment analysis in panel F is based on hypergeometric test.

We evaluated the ovarian metastatic samples using PCA and annotated them by histological subtype, PAM50, *CDH1* expression and ESTIMATE purity scores (**Figure 3C**, **Supplementary Data 2**). While there was no clear separation by histology and purity, we did see an obvious separation by PAM50. In addition, *CDH1* low and *CDH1* high metastatic samples clustered distinctly. As expected, there was a correlation between *CDH1* expression in the metastatic samples and histological subtype, with ILC metastases expressing significantly less *CDH1* compared to IDC metastases (**Figure 3D**). To further explore potential subtype-associated differences in ovarian metastasis, we performed differential expression analysis between ILC and IDC metastases. We found 35 significant differentially expressed genes (DEGs) between ILC and IDC (**Figure 3E**; DEG list in **Supplementary Data 3**). DEGs with higher expression in IDC included *CDH1, LHX8, KRT5, MEIOC, PPP1R14C, ATG9B, TUBB3, CDH18* and *ZIC5*, and those with higher expression in ILC included *CYP2B6*, *BPIFB2, MYBPC1, UCP1, PGLYRP2* and *PCSK1N* among others. The two top upregulated genes in ILC ovarian metastases *(CYP2B6, BPFIB2)* have roles in lipid (and steroid) metabolism [39], but the limited number of DEG does not allow further pathway analysis.

### Transcriptomics identifies overexpression of G-protein coupled receptors in ovarian metastasis

To identify DEGs between primary tumors and ovarian metastases, we corrected for potential background contamination from normal ovarian tissue (for details see **Supplementary Figure S6**, **Supplementary Data 4, 5, 6** and Methods). Background corrected DEGs are visualized using a volcano plot in **Figure 4A** and a heatmap in **Figure 4B** (complete list of DEGs can be found in **Supplementary Data 6**). Ingenuity Pathway Analysis (IPA) analysis revealed glutamate receptor (GluR) signaling (geneset comprised of following genes: *GRIA4*, *GRIK2*, *GRIK4*, *GRIN3A*, *GRM7*, *GRM8* and *SLC17A7*) as the most up-regulated pathway in the ovarian metastases (**Figure 4C**; detailed list in **Supplementary Data 8**). Given the previously described role of glutamate signaling in breast cancer metastasis [40] and in endocrine resistance in ILC [41, 42], we further assessed its activity in the context of ovarian metastasis. Briefly, glutamate receptors (GluR) can be divided into ionotropic GluR (iGluR) and metabotropic GluR (mGluR) with the former being ion channel associated receptors and the latter being GPCRs. Both metabotropic and kainite GluR signaling showed significantly higher activation (based on GSVA scores) in ovarian metastases compared to primary tumors (**Figure 4D**; **Supplementary Figure S7A**). On the other hand, iGluR (NMDA, and AMPA) were upregulated in brain metastases (**Supplementary Figure S7B**) as previously reported [40] but not in the ovarian metastases. In contrast, the kainate receptor *GRIK2* was significantly upregulated in ovarian metastases, although this was not specific for the ovarian metastases, as we observed similar upregulation in brain metastases (**Supplementary Figure S7C**). And finally, a number of the mGluR GPCRs (*GRM7* and *GRM8*) were significantly upregulated in ovarian metastases and upregulation of *GRM7* was specific to ovarian metastases (**Supplementary Figure S7C**).

**Figure 4.**
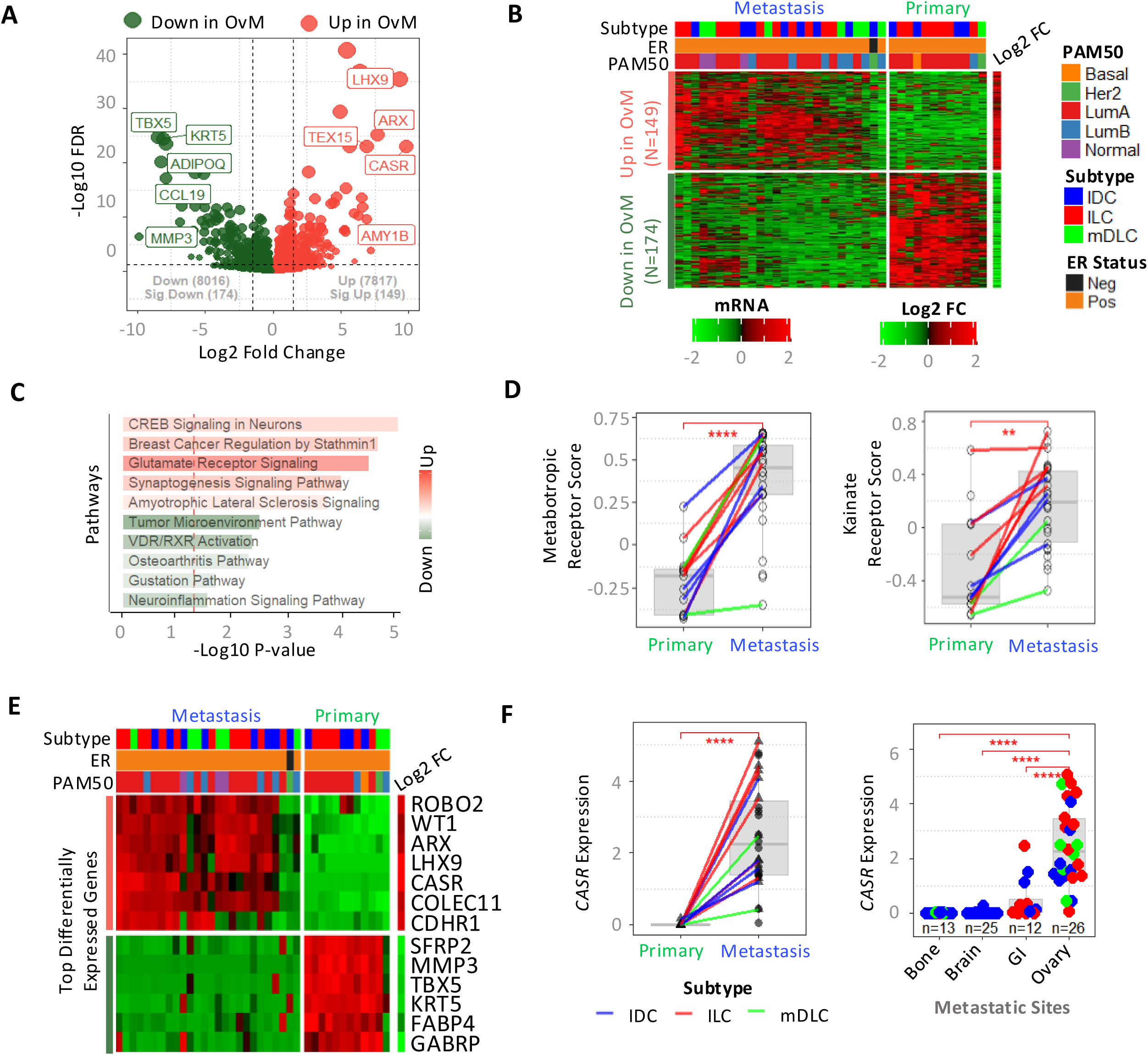
Identification of potential drivers of ovarian metastasis. (A) Volcano plot of DEGs associated with ovarian metastasis (OvM) after background correction (see Sup. Fig. 6). (B) Heatmap highlighting background corrected ovary metastasis up- and down-regulated DEGs. (C) IPA pathway analysis top hits based on adjusted p-value. Color bar shows activation z-score values, where red shows positive scores (up-regulated in ovary metastasis) and green shows negative scores (down-regulated in ovary metastasis). Only top 5 up-regulated (+ive activation z-score) and down-regulated (-ive activation z-score) pathways are shown. (D) Paired plots showing GSVA scores between primary and metastatic samples for glutamate receptor genesets. P value based on Wilcoxon signed rank test. Dots without line represent orphan metastasis samples. (E) Heatmap showing top DEGs (including *CASR*) based on effect size cutoffs of 2SD from mean effect size. (F) *CASR* expression changes from primary breast tumor to ovary metastasis (left) and to various other site metastasis (right) in breastMETs dataset (see “Additional RNAseq datasets” section in methods).

To extend our search for potential drivers of ovarian metastasis, we next focused on DEGs with effect size greater or less than 2 standard deviations from the mean effect size of all significant DEGs. This identified 7 up-regulated DEGs: *ROBO2, CASR, COLEC11, ARX, LXH9, WT1* and *CDHR1* and 6 down-regulated DEGs: *FABP4, SFRP2, MMP3, TBX5, KRT5* and *GABRP* (**Figure 4E**). Similar to the mGluR, the Calcium-Sensing Receptor CaSR is a member of the GPCR family [22, 43, 44]. Up-regulation of CaSR was most noticeable in ILC metastatic pairs, although up-regulation was also observed in both IDC and mDLC metastatic pairs (**Figure 4F**, left). The difference in CaSR mRNA expression between ILC and IDC metastases did not reach statistical significance (wilcox.test p-value 0.12). CaSR expression was significantly higher in ovarian metastasis than other (bone, brain, and GI) sites of metastasis (**Figure 4F**, right). CaSR displayed higher expression in normal ovary compared to normal breast tissue (**Supplementary Figure S7B**), however the up-regulation seen in the ovarian metastases was of greater magnitude and passed background correction (**Supplementary Figure S6A & B**). In summary, RNAseq analysis identified a potential role for GPCRs, and specifically a number of mGluR and CaSR in breast cancer metastases to the ovary. Previous studies have examined the role of CaSR in breast cancer bone metastasis [45–47], but not in the context of ILC or in ovarian metastasis.

### Activated CaSR promotes migration and F-actin polymerization in ILC cell lines via MEK/ERK pathway activation

CaSR is highly expressed in the parathyroid glands and kidneys, where it maintains calcium (CaCl_2_) homeostasis through regulating parathyroid hormone secretion and subsequent bone and urinary CaCl_2_ reabsorption [48]. To understand its role in ILC, we overexpressed CaSR (CaSR-OE) in four ILC cell lines: MDA-MB-134, BCK4, MDA-MB-330 and SUM44 (**Figure 5A**; **Supplementary Figure S8A**) and stimulated its activity with CaCl_2_ or the calcimimetic R568. Configured CaSR did not stimulate proliferation (data not shown).

**Figure 5.**
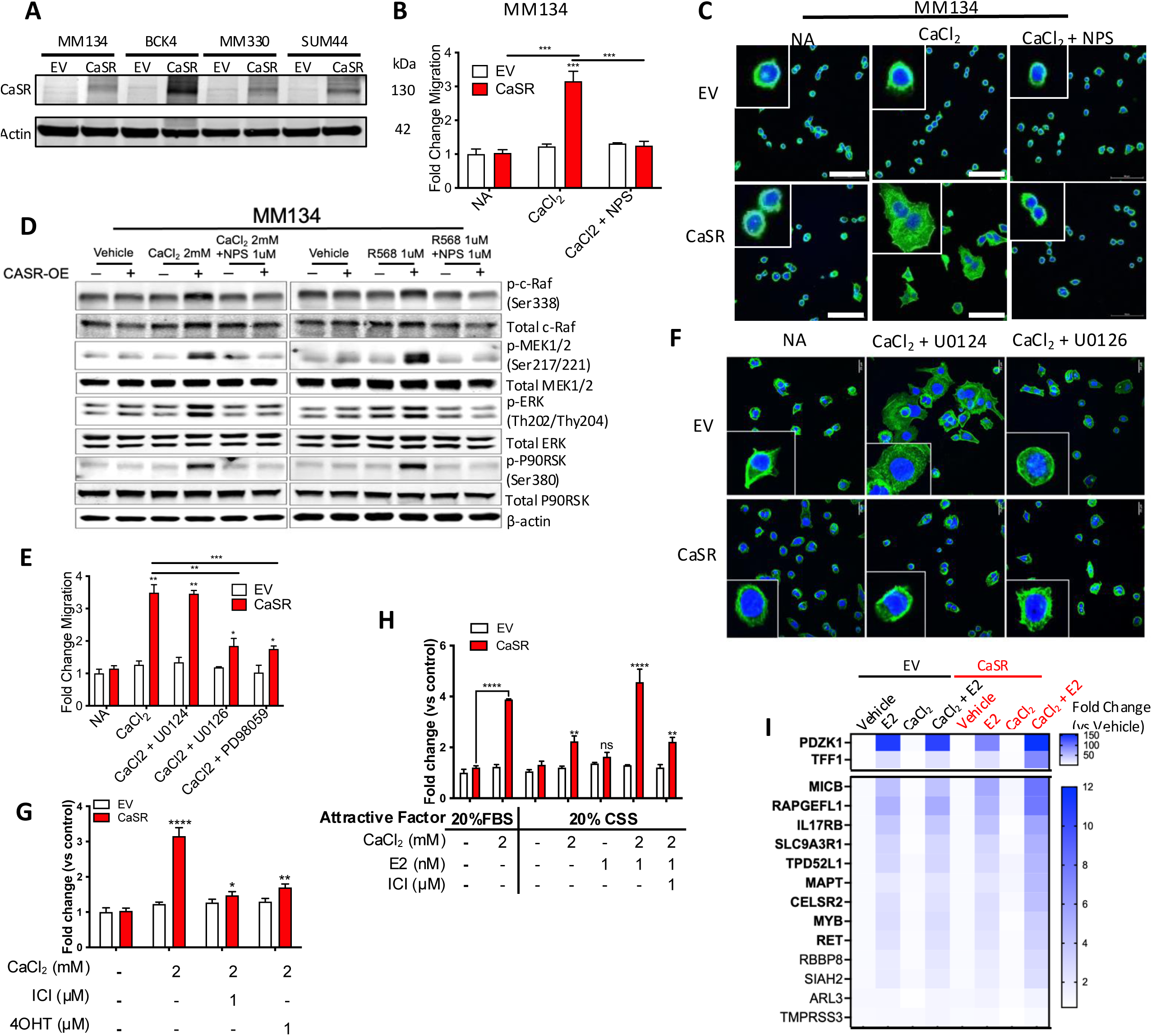
Activated CaSR promotes MEK/ERK-dependent migration and F- actin reorganization in ILC cell lines, enhanced by estrogen. (A) Western Blotting using the indicated antibodies on whole cell lysates from CaSR-OE and empty vector control (EV) of MDA-MB-134 (MM134), BCK4, MDA-MB-330 (MM330), and SUM44. β-actin was used as loading control. (B) Quantifications of crystal violet-stained inserts with CaSR-OE and EV MM134 from chemotaxis toward FBS after 72 hours. Migration toward FBS is induced by CaCl_2_ (2mM) in CaSR-OE cells and abrogated by treatment with the calcilytic NPS (1µm). Data representative of at least 2 experiments (n=3). P-values are from unpaired t-tests. ***, *P* ≤ 0.005. (C) Confocal microscopy imaging of F-actin staining (green) in CaSR-OE and EV MDA-MB-134. CaCl_2_ treatment enhanced F-actin spreading per cell while NPS abrogated the effect. Hoechst staining (blue) was used for nuclear staining. 50µm scale bar. Each experiment has been repeated at least 2 times (n=3). (D) Western Blotting using the indicated antibodies on whole cell lysate from CaSR-OE and EV MDA-MB- 134 cells treated without or with CaCl_2_ (2mM) or R568 and NPS (1µM) for 1 hour. (E) quantification of crystal violet-stained inserts with CaSR-OE or EV MDA-MB-134 from chemotaxis toward FBS after 72 hours with treatments of MEK inhibitors (U0126 and its negative control U0124 (1 μM) and PD98059 (5 μM)). MEK inhibition prevents CaCl_2_ induced chemotaxis toward FBS in CaSR-OE MDA-MB-134. Data representative of 2 experiments (n=3). P-values are from unpaired t-tests. *, *P* ≤ 0.05. **, *P* ≤ 0.005. ***, *P* ≤ 0.0001. (F) Confocal microscopy imaging of F-actin staining (green) of MDA-MB-134 with and without MEK inhibitors. (G), ER inhibitors (fulvestrant (ICI) and tamoxifen (4OH)) prevents CaCl_2_ induced chemotaxis toward FBS in CaSR-OE MDA-MB-134. (H) In hormone deprived conditions, estradiol (E2) enhances CaCl_2_ induced chemotaxis toward FBS in CaSR-OE MDA-MB-134 which is suppressed with fulvestrant treatment. Both (G, H) are representative of at least 2 experiments (n=3). P-values are from Dunnett’s multiple comparison test or t-test comparing each group to CaSR-OE with no treatment **, *P* ≤ 0.05. ***, *P* ≤ 0.005. ****, *P* ≤ 0.0001. (I) Heatmap illustrates the relative mRNA expression of the 15 consistently intersected E2 regulated genes with CaSR activation. *CASR*-OE and EV MDA-MB-134 cells were treated with 1nM E2, 2 mM CaCl2 or their combination for 24 hours. qRT-PCR was performed, and gene expressional fold changes were calculated by ΔΔCt method normalized to their own vehicle. Mean fold changes (n=3) are shown by the color key.

Activated CaSR was previously shown to promote migration [49], and we therefore examined if CaSR overexpression increased serum-induced migration of ILC cells which in general show low migration and chemotaxis [50]. Activated CaSR enhanced migration across tested ILC lines (**Figure 5B**; **Supplementary Figure S8B-D**) and was reproduced with the calcimimetic R568, although less than with CaCl_2_ (**Supplementary Figure S8E**, **S8F**). These effects could be blocked with the CaSR antagonist NPS2143, which was used at a dose that did not induce cell death (**Supplementary Figure S8G**). We confirmed the effect of activated CaSR on migration with a wound-scratch assay (data not shown) but found no promotion of invasion in a transwell chamber coated with collagen I or Matrigel (**supplementary Figure S8H**).

Activated CaSR triggers filamentous actin (F-actin) polymerization in human embryonic kidney (HEK)-293 cells [51], a process required for cell migration [52]. Phalloidin staining of MDA-MB-134 cells revealed increased F-actin staining in activated CaSR-OE cells, which also showed enlarged cytoplasm and increased protrusions compared to empty vector cells (**Figure 5C**). Treatment with NPS2143 abrogated these effects, confirming their CaSR-dependency. Together, our results suggest that activated CaSR is sufficient to promote migration in ILC cell lines, without activating proliferation or invasion, and that this is likely mediated at least in part via CaSR’s role in actin polymerization.

The MEK/ERK pathway regulates cell motility and has been shown to be activated by CaSR in breast cancer cell lines [49, 53]. We confirmed this in MDA-MB-134 cells where we observed increased phosphorylation of c-Raf, MEK1/2, ERK and P90RSK in CaSR-OE cells in the presence of CaCl2 or R568, which was blocked by NPS2143 (**Figure 5D**). MEK/ERK activation was detected as fast as 10 min after CaCl_2_ supplementation and lasted for at least 6 hours (**Supplementary Figure S9A**).

To determine if inhibition of the MEK/ERK pathway could reverse CaSR-mediated effects on migration, we utilized MEK1/2 inhibitors PD98059 and U0126 [54, 55], along with the inactive compound U0124 as a negative control. Dose response experiments in MDA- MB-134 determined that 5μM PD98059 and 1μM U0126 were non-toxic doses (**Supplementary Figure S9B**) and effectively blocked the MEK/ERK pathways after 1 hour without affecting growth over 3 days of treatment (**Supplementary Figure S9C**). Transwell migration experiments revealed that PD98059 and U0126 blocked the increase in migration seen in CaCl_2_ activated MDA-MB-134 CaSR-OE cells, whereas U0124 failed to exert an effect (**Figure 5E**; **images in S9D**). Furthermore, addition of U0126 to CaCl_2_ activated MDA- MB-134 CaSR-OE cells reduced the F-actin spreading and overall F-actin staining per cell, whereas again the addition of U0124 had no effect (**Figure 5F**; **quantification in S9E**).

Altogether, our findings indicated that CaSR activates the MEK/ERK pathway, which in turn modulates the actin cytoskeleton to increase migration.

### Activated CaSR requires functional ER alpha for migration

Previous studies have shown that activated CaSR enhances ER activity in MCF7 cells [56], and we therefore investigated whether activation of CaSR in ovarian metastases was associated with altered ER signaling. We divided the ovarian metastatic cohort into high (n=12) and low (n=13) *CASR* expression groups and performed gene signature enrichment analysis. We found an enrichment for early and late estrogen response in the high *CASR* expressing group, suggesting higher ER activity in these tumors (**Supplementary Figure S10A**). To then test whether activated CaSR-induced migration is dependent upon ER activation, we repeated the assays in the presence of the ER inhibitors 4-OH-tamoxifen (tam) and fulvestrant (ICI). Both ER inhibitors abrogated activated CaSR-induced migration (**Figure 5G**; **Supplementary Figure S10B**).

To further confirm the role of estrogen in CaSR-induced migration, cells were deprived in phenol red-free and charcoal-stripped bovine serum (CSS) conditions, in which the addition of CaCl2 induced a 2.2-fold migration, while there was no significant effect of estradiol (E2) alone. However, the combination of CaCl2 and E2 increased migration to 4.5- fold, indicating that estrogen enhances the migration response elicited by activated CaSR. Treatment with ICI blocked the E2/CaCl2-mediated migration (**Figure 5H**; **Supplementary Figure S10C**).

To identify specific CaSR-associated ER downstream genes, we overlapped the core- enriched estrogen response signature genes in *CASR*-high ovarian metastatic tumors (n=12) and in E2-induced MCF7 (GSE89888), T47D (GSE89888) and MDA-MB-134 (GSE50695) cell lines which revealed 15 estrogen responsive genes (**Supplementary Figure S10D**). 11 of these genes exhibited a stronger E2 response in CaCl_2_ stimulated CaSR-OE cells compared to EV cells (**Figure 5I**) by qPCR, confirming that activated CaSR enhances ER activity. Taken together, our data suggests interaction between activated CaSR and ER signaling at the level of transcriptional regulation and induction of migration in ILC cells.

## Discussion

Metastatic breast cancer is a major public health concern as it accounts for greater than 90% of breast cancer-related deaths [57]. Over the past decade, increased access to metastatic samples has allowed comprehensive studies discerning patterns of metastatic spread according to breast cancer subtype. Here we took a step forward in understanding molecular features of breast cancer that contribute to metastasis to the ovary. This is of particular importance to further understand the invasive lobular histological subtype of breast cancer, which is afflicted by ovarian metastasis three times as often as the invasive ductal histological subtype [10].

Our analysis of the Foundation Medicine cohort is the largest genomic characterization of breast cancer ovarian metastases reported to date. The enrichment of *CDH1*, *PIK3CA,* and *TBX3* mutations and attenuation of *TP53*, *ESR1,* and *MYC* mutations in ovarian metastases support the association between ovarian metastasis and ILC-like genomic features. However, only 47% of ovarian metastases harbored *CDH1* mutations, and 16q arm- level loss was reduced relative to local ILC (64% vs 84%), indicating that some breast cancer ovarian metastases arise from non-ILC primary tumors. This is reflected in our UPMC cohort, where 30% (8/27) of cases were IDC. We also observed a lower frequency of HRDsig- positive cases in ovarian metastases (8.1%) compared to non-ovarian metastases (14.3%), and in local ILC (6.6%) compared to local breast cancer overall (17.6%), suggesting that homologous recombination deficiency is not a major driver of breast cancer metastasis to the ovary.

Our UPMC cohort of matched primary and metastatic samples is reflective of known features of breast metastasis to the ovaries including enrichment of ILC, younger median age (43 years), and largely pre-menopausal status [17]. Kutasovic et al. recently reported the largest study to date comparing 54 breast cancer patients with gynecologic metastasis to 455 patients without gynecologic metastasis, including 15 cases of matched primary and metastatic tumors. In line with the findings presented here, they found that patients with gynecologic metastasis were younger, pre-menopausal women with ILC and luminal breast cancer [58]. Our cohort faithfully recapitulated E-cadherin and p120 staining of their respective subtypes as well (**Figure 2B**). Mutation analysis revealed *PIK3CA, CDH1, KMT2C, FOXA1,* and *RUNX1* as the top 5 mutated genes (**Figure 2D**), several of which were identified by Desmedt et al. as key mutations in ILC [6] and enriched in the metastatic versus primary tumors in gynecologic metastases cohort reported by Kustasovic et al. [58]. We found mutations enriched in ovarian metastases compared to the primary tumors including *ERBB3, KMT2C, NRAS, JAK3,* and *AKT1*. Most of these mutations have been previously described to enriched in breast cancer metastases [59–61], and might not be specific drivers for metastases to the ovary but a reflection of activation of RTK and TF pathways in endocrine resistance as previously described [60]. For example, *ERBB3* mutations can activate *ERBB2*/*ERBB3* signaling resulting in endocrine resistance [62, 63]. Similarly, alterations in PI3K/Akt signaling are known drivers of resistance to hormonal therapy [64]. An exception is the *JAK3* V722I which has not previously been identified in breast cancer, and in general *JAK3* mutations are mostly found in hematological malignancies [65, 66]. Of note, the MammaSeq panel used for mutation profiling is limited to a curated set of known actionable mutations in select breast cancer genes. Hence, our analysis does not provide a comprehensive overview of all possible mutations and expanding the study with additional samples and WGS is planned for the future. We did also study CNV and identified frequent amplifications of *FGF19*, *FADD* and *CTTN*, all located in 11q13.3, a region that is well known to be amplified in a number of solid tumors including breast cancer [67].

Our transcriptomic analysis and PAM50 subtyping coupled with principal component analysis did not identify a unique molecular feature of lobular metastasis to the ovary in our cohort. IDC and ILC metastatic samples clustered together, although there was a trend towards separation based on *CDH1* expression (**Figure 3C**). Molecular subtype was better able to separate the samples. The lack of clear separation solely based on histological subtype is also reflected in the small number of DEGs between ILC and IDC metastases (**Figure 3E**). This was somewhat surprising, and could have a number of reasons, including limited sample size, known problems with diagnosis of ILC [68], or a lobular-like phenotype in the IDCs that metastasize to the ovary.

When comparing primary tumors and ovarian metastases (**Figure 4B**), pathway analysis identified the GluR signaling pathway as the most upregulated pathway in the metastases (**Figure 4C**). Previous analyses have identified roles for mGluR in tamoxifen resistant ILC cells [41], and in patients treated with tamoxifen [69]. Intriguingly, there was some organ specificity, as our analyses validated previously reported upregulation of the ionotropic GluR NMDAR and AMPA in the brain [40, 70] but not in the ovary. We identified ovarian-specific upregulation of *GRM7* and *GRM8*, two GPCRs deserve further study as potential drivers of metastases.

Analysis of the top-upregulated genes in the ovarian metastases identified another GPCR, the calcium sensing receptor CaSR [44] (**Figure 4E**). It was upregulated in both ILC and IDC, and to a greater extent in ILC although this was not significant. CaSR expression was significantly higher in metastases to the ovary compared to other sites of metastases (**Figure 4F**).

We performed functional studies on CaSR in ILC cell lines and found that CaSR activation increased migration which could be blocked with CaSR inhibitor (**Figure 5A-E**).

These results agree with previous studies showing that CaSR activation increases migration of IDC cell lines [49]. ILC cell lines show very limited migratory ability [50] and thus our results emphasize a role of CaSR in this part of the metastatic process. The lack of CaSR to mediate effects on invasion suggest that other pathways synergize with its activity in the complex process of distant metastases. Downstream mechanistic analysis revealed that activated CaSR increases F-actin formation (**Figure 5C**) through MEK/ERK signaling activation (**Figure 5F**), a well characterized downstream pathway of CaSR [49, 53, 71]. A recent study identified a role for CaSR in cytokine secretion [72] which could support tumor progression.

We also discovered a strong association between ER signaling and CaSR expression in the ovarian metastases, and the effects of CaSR on migration were dependent on ER (**Figure 5G-I**). This novel finding complements previous literature showing in the context of bone metastases that calcium released from the lesions was able to modulate ER through CaSR activity [56]. With 44, the average age of patients at the time of diagnosis in our cohort is substantially lower than that of the average breast cancer patient, and many patients are likely to have received tamoxifen as part of their adjuvant treatment. In addition, it is known that there can be a residual low E2 level in pre-menopausal breast cancer patients treated with endocrine therapy [73]. It is therefore tempting to speculate that residual estrogen in the ovaries might provide a favorable environment for the breast cancer cells.

Another unanswered question is how CaSR is activated in tumors? ER+ ILC and IDC are highly metastatic to bone, and it is plausible that patients in our cohort presented with simultaneous bone metastasis. Notably, Kutasovic et al. found that 45.8% of patients in their gynecologic metastases cohort presented with metastases to the bone [58]. Bone metastasis is known to induce osteolysis, which leads to extensive calcium release [56] and could potentially activate CaSR. Whether calcium release from bones directly promotes ovarian metastasis remains to be elucidated.

In summary, we have identified candidate drivers of ovarian metastasis including clinically actionable mutations such as *ERBB3, JAK3* and *PIK3CA*, and glutamate receptor pathway activation. Our comprehensive multi-omics analysis revealed CaSR as a promising mediator of ILC migration, however, other GPCRs like glutamate receptors such as *GRM7* and *GRM8* deserve further exploration for their role in ovarian metastases. Larger functional genomics screens would be appropriate to dissect their role in diverse steps of ovarian metastases. And finally, we hope that the comprehensive data set presented as part of our study will serve as a valuable resource to those in the metastatic breast cancer research community studying ovarian metastasis and lobular breast cancer.

## Methods

### Medicine cohort genomic analysis

Mutations and copy number changes were analyzed from 15,613 local breast cancer tissues (of which 1,350 were from patients with ILC), 22,010 non-ovarian metastases, and 246 ovarian metastases, all of whom underwent tumor-only sequencing using the FoundationOne (F1Cdx and F1) assays. An investigational scar-based measure of HRD (HRDsig), incorporating copy number and indel features, was called using a machine learning-based algorithm and offered as a laboratory professional service.

### UPMC Cohort Sample acquisition

Collection of 49 samples (12 primary, 27 metastases, 6 normal breast, 4 normal ovaries) was performed in collaboration with Pitt Biospecimen Core (PBC) and pathologists at UPMC Magee Women’s Hospital under Institutional Review Board (IRB) guidelines. 13 ovarian metastases were matched with their 12 primary tumors (two metastases Ov11 M1 & M2 shared the same primary Ov11 P), while 14 metastases (two metastasis Ov14 M1 & M2 were from same patient) were unmatched (“orphan” samples). Clinicopathological features of the patients/tumors are illustrated in Table 2 and **Supplementary Table S1**. In addition, 6 normal breast and 4 normal ovarian samples were collected for comparison and quality control in the subsequent analyses. Analysis of histological classification was performed by two pathologists (PL and EE).

### FFPE processing and nucleic acid isolation

Formalin-fixed paraffin-embedded (FFPE) samples were macro-dissected for RNA and DNA isolation (3 to 6 sections per sample depending on tumor size and cellularity) using the Qiagen AllPrep FFPE kit as per manufacturer’s instructions. RNA and DNA concentrations were determined with the Qubit 3.0 Fluorometer (ThermoFisher Scientific). RNA fragment sizes distributions (DV200 metrics) were obtained utilizing either the Agilent 2100 Bioanalyzer or the Agilent 4200 TapeStation. Distribution of the samples based on tumor cellularity is shown in Table S2.

### RNA sequencing, Purity Analysis, PAM50, Differential Expression Analysis, Pathway analysis

See supplementary methods

### Mutation Analysis and Nanostring Copy Number Analysis

See supplementary methods

### Droplet Digital PCR (ddPCR)

Bio-Rad QX100 droplet digital PCR platform was used for sensitive detection of *CDH1* mutations in patient ov11 which had 2 ovarian metastases. Primers and probes were ordered for R74* (Integrated DNA Technologies) mutations. Forward primer sequence was: 5’- GGTCGACAAAGGACAGCCTATTTT-3’, and reverse primer sequence was: 5’- GGCCTTTTGACTGTAATCACACCAT-3’. 60ng of control samples were processed for ddPCR analysis as previously described [74]. Briefly, DNA samples were combined with primers, probes, and supermix, and then added to cartridge. Droplets were generated with the droplet generator and transferred into a 96-well plate for PCR amplification, and a droplet reader was used to count PCR positive and PCR negative droplets.

Data were analyzed using QuantaSoft^TM^ packages (BioRad) where target concentration was normalized to reference concentration and multiplied by the number of reference loci in the genome to generate copy number calls.

### Cells and reagents

See supplementary methods ***RNA extraction and qPCR*** See supplementary methods

### Immunoblotting

See supplementary methods

### Proliferation, migration, transwell chemotaxis and haptotaxis, and invasion assays

See supplementary methods

### Hormone Deprivation

See supplementary methods

### Immunofluorescence (IF) staining

The cells were grown on coverslips and fixed with ice-cold 4% paraformaldehyde for 10 min. After a PBS wash, the cells were blocked for 30 min with filtered 5% BSA and 0.3% triton X in PBS. The cells were then incubated with primary antibodies against CaSR (1:1000 dilution; Sigma-Aldrich) overnight at 4 °C. Lastly the coverslip was incubated with Cy5-IgG fluorescence secondary antibody (1:200 dilution; Invitrogen) and Hoechst for DAPI (1:10000 dilution; Thermo Fisher Scientific) for 1h at room temperature. Then, the images were taken with Nikon A1 advanced confocal. The images were then processed with ImageJ.

#### Immunohistochemistry (IHC)

Cell pellets were fixed in 10% formalin and processed for paraffin embedding. Antigen retrieval was performed using citrate buffer and stained with CaSR antibody (1:1000 dilution; Sigma-Aldrich) overnight at 4 °C. Endogenous peroxidases were blocked using 0.3% H_2_O_2_ in TBS for 15 minutes. Secondary antibody (DAKO) was used followed by 3,3’- diaminobenzidine (DAB, DAKO) and then counterstaining with hematoxylin (Sigma-Aldrich).

#### F-actin staining

The cells were grown onto coverslips then fixed with 4% paraformaldehyde for 10 min at room temperature. Permeabilization was done with 0.1% Triton X-100 for 5min and then stained for F-actin and the nucleus with the fluorescent Alexa Fluor 488 Phalloidin (Thermo Fisher Scientific) and Hoechst (1:10000 dilution; Thermo Fisher Scientific) for 20 min at room temperature. The coverslips were washed with PBS and mounted onto slides. Images were taken with Nikon A1 advanced confocal. The images were then processed with ImageJ. The fluorescence intensity of the green channel was divided by the number of nuclei per field which yielded the fluorescence intensity per cells. 5 fields of each triplicate were measured.

#### Selecting high confident CASR-related ER Target Genes

Three estrogen treatment data sets were downloaded from GSE89888 (MCF7 and T47D) and GSE50695 (MDA-MB-134) [77, 78]. GSEA on “Estrogen Response Early” and “Estrogen Response Late” hallmark signatures were performed between vehicle and E2 groups of each data sets. Core enriched genes derived from early and late response signatures were further intersected for each data set to obtain the cell line specific E2 regulated gene list. For *CASR*- regulated ER targeted genes, core enriched genes between *CASR* high and low ovary metastatic tumor groups derived from early and late response signatures GSEA analysis were intersected. Cell line specific E2 regulated genes from MCF7, T47D and MDA-MB-134 overlapped with *CASR*-regulated ER targeted genes to obtain the 15 highly confident *CASR*- related ER target gene list. Venn Diagram was generated with https://bioinformatics.psb.ugent.be/webtools/Venn/.

### Statistical analysis

Statistical analysis was performed using GraphPad Prism 7 (GraphPad Software Inc., La Jolla, CA). Data are presented as mean and ± SD of 1 of the technical triplicates of 1 of the biological triplicates. Statistical tests used for each figure are indicated in the respective figure legend. For growth curves and wound-scratch migration assays, two-way ANOVA was performed. For transwell chemotaxis assays and quantification of fluorescence staining unpaired t-test were performed. For quantification of the qRT-PCR, gene expressional fold changes were calculated by ΔΔCt method normalized to their own vehicle.

## Author’s Contribution

**Designing Research Studies:** Y. Qin, A. Basudan, L. Savariau, O. Shah, L Coffman, A.V Lee, S. Oesterreich

**Collecting and characterization of sample, and conducting experiments:** Y. Qin, L. Savariau, O. Shah, A. Basudan, E.Elishaev, P. Tallapaneni

**Analyzing Data:** Y. Qin, L. Savariau, O. Shah, A. Basudan, Z. Li, T. Liu, N. Tasdemir, P. Lucas, A.V Lee, J.M Atkinson, S. Oesterreich

**Writing, review, and/or revision of the manuscript:** C. Merkel, L. Savariau, O. Shah, Y. Qin, O. McGinn, L. Coffman, E. Elishaev, J.M Atkinson, P. Lucas, A.V Lee, S. Oesterreich **Study supervision:** J.M Atkinson, A.V Lee, S. Oesterreich

**Method authorship order among co-first authors:** Functional studies which uncovered key phenotypes of CaSR manipulation in breast cancer cells were initiated by Y.Qin and validated and completed by L.Savariau. A. Basudan initiated tissue collection and processing and performed initial transcriptomic analysis identifying CaSR as a potential target to be investigated. O. Shah extended and finalized bioinformatic work to comprehensively characterize the cohort (including methodology for background correction), highlighting GPCR pathway up-regulation in ovarian metastases. The majority of the final manuscript was drafted by L. Savariau and O. Shah and edited by authors indicated above. The four authors provided equal intellectual contributions to the research study.

## Acknowledgments

This project used the University of Pittsburgh Health Sciences Core Research Facilities Genomics Research Core (<u>RRID: SCR_018301</u>), the UPMC Hillman Cancer Center and Tissue and Research Pathology/Pitt Biospecimen Core shared resource which is supported in part by award P30CA047904. This study used high-performance research computing core (RRID: SCR_022735) at the University of Pittsburgh Center for Research Computing supported by NIH S10OD028483 (to AVL). The authors would like to thank Jian Chen for technical assistance, Dr Nolan Priedigkeit for bioinformatics input, Dr John Wysolmerski for advice on CaSR, Dr Wenhan Chang for help with our efforts for CasR IHC, Dr Rebecca Riggins for some early discussions on the role of glutamate receptors in ILC, and Dr Aatur Singhi for providing controls for IHC. Research funding for this project was provided in part by Susan G. Komen Scholar awards (SAC110021 to AVL and SAC160073 to SO), the Breast Cancer Research Foundation (to AVL and SO), the Magee-Womens Research Institute and Foundation, The Shear Family Foundation, The Metastatic Breast Cancer Network and the Nicole Meloche Foundation. ABS was supported in part through the Vice Deanship of Scientific Research Chairs; Research Chair of Medical and Molecular Genetics at King Saud University (2014-2018).

## Supplementary materials

### Supplementary Figure Legends

**Supplementary Figure 1. Association between ovarian metastasis and histological subtype**

Association between metastatic sites [ovary and other (bone, brain & GI)] and histological subtype (ILC, IDC & mDLC) in breastMETs cohort was tested using Pearson’s chi-square test. Residuals reflect significant enrichment (blue) or depletion (red). P-value reflects significant association. Standardized Pearson residuals are defined as the differences between observed and expected values divided by square root of expected value and p-value was computed from a chi-squared distribution. Two ILC patients Ov11 and Ov14 had multiple metastatic samples. For each of them only a single sample was considered for this association analysis.

**Supplementary Figure 2: MammaSeq^TM^ based DNAseq QC**

(A) Probe coverage in the two pools of the MammaSeq panel across profiled samples. (B) Mean coverages across profiled samples. (C) SNV count before and after filtering. M = Metastasis and P = Primary Tumor.

**Supplementary Figure 3. Mutational landscape of all (paired + orphan) ovarian metastatic samples and clinical actionability analysis**

(A) Genes with largest mutational frequency change between primary and all metastatic samples. (B) Oncoprint showing top 20 altered genes in all profiled samples. Samples are split by tumor subtype i.e., IDC, ILC and mDLC and tissue type i.e., primary and metastasis. (C) Clinically actionable mutations identified in the MammaSeq profiled ovary metastasis cohort. (D) Oncoprint showing only clinically actionable alterations across samples.

**Supplementary Figure 4. Ov11 *CDH1* R74* mutation assessment in RNA/DNA reads**

(A) *CDH1* gene coverage in MammaSeq panel. (B) *CDH1* R74* (C>T) mutation at chr16:68,835,629 visualized using IGV for RNAseq and DNAseq (MammaSeq) reads from respective bam files.

**Supplementary Figure 5. RNAseq QC and exploratory analysis**

(A) QC assessment of RNAseq reads and identification of samples that fail mapped read QC.

(B) Genotype similarity analysis to confirm matched status of samples from same patient using SNPs in RNAseq reads. (C) Sample-sample similarity based on Euclidean distance. Top 10% variable genes were used. (D) PCA using top 10% variable genes (using DESeq2 VST normalized counts) after removal of Ov3M. Samples annotated by tissue type, PAM50, subtype and ESTIMATE purity scores

**Supplementary Figure 6. DEA scheme for identification of ovarian metastasis associated DEG**

(A) Differential expression analyses (DEA) scheme used to identify ovary metastasis differentially expressed genes (DEGs) and correction of potential background due to contamination from normal ovary (nOvary). Contamination of Metastasis Up – conMet Up and Contamination of Metastasis Down – conMet Dn.(B) Up DEG background correction – cutoff of 0.5 log2FC used to classify DEGs as true signal (up in ovary met) or background contamination (up in ovary site). (C) conMet Dn DEG background correction – cutoff of 0.5 log2FC used to classify DEGs as true signal (down in ovary met) or background contamination (down in ovary site). LFC = Log2FC.

**Supplementary Figure 7. Glutamate receptor family signaling activity in ovary, bone, GI and brain metastasis**

(A) Heatmap of GSVA scores for glutamate receptor genesets across ovarian metastatic samples and paired primary breast tumors. (B) Paired plots of GSVA scores for glutamate receptor genesets across bone, brain and GI metastasis and corresponding paired primary breast tumors. (C) Paired plots of *GRIK2*, *GRM7* and *GRM8* mRNA expression across bone, brain, GI and ovary metastasis and corresponding paired primary breast tumors.

**Supplementary Figure 8. CaSR overexpression and calcimimetic activation promote migration and growth in ILC cell lines, reversed by calcilytic inhibition**

(A) Confocal microscopy of CaSR (pink) in CaSR-OE and EV ILC cell lines show localization at the membrane and cytoplasm. DAPI staining (blue) was used for nuclear staining. Scale bars 50 µm. (B) Images of crystal violet-stained inserts with CaSR-OE and EV ILC cell lines from chemotaxis toward FBS after 72 hours. Quantification shown in Figure 5B. (C&D) Images (left) and quantifications (right) of crystal violet-stained inserts with CaSR-OE and EV ILC cell lines from chemotaxis toward FBS after 72 hours. Migration toward FBS is induced by CaCl_2_ (2mM) or the calcimimetic R568 (5µm) in CaSR-OE cells and abrogated by treatment with the calcilytic NPS (1µm). Data representative of at least 2 experiments (n=3). P-values are from unpaired t-tests. ***, *P* ≤ 0.005, ****, *P* ≤ 0.001. (E, F) Images and quantifications (bottom) of crystal violet-stained inserts with CaSR-OE and EV ILC cell lines with indicated treatment from chemotaxis toward FBS. (E&F) The calcimimetic R568 induce migration in CaSR-OE MDA-MB134 and MDA-MB-330. P-values are from t-test analysis. ***, *P* ≤ 0.005. (G) Proliferation curves of CaSR and EV MDA-MB-134 cells with and without CaCl_2_ treatment and with NPS2143 or R568 for 3 days. Arrows indicate the drug concentration chosen for further experiments. (H) Images of crystal violet-stained inserts with CaSR-OE and EV MDA-MB-134 from Collagen I and Matrigel invasion assays. Activated CaSR by CaCl_2_ did not improve invasion to either ECM. MDA-MB-231 cells were included as a positive control to confirm that each matrix (Collagen I and Matrigel) supported invasion under the assay conditions used.

**Supplementary Figure 9. Activation of CaSR induces MEK/ERK signaling and can be blocked with calcilytic**

(A) Western Blotting with indicated antibodies on whole cell lysate from CaSR-OE and EV MDA-MB-134 treated with CaCl2 with indicated duration. (B) Proliferation curve of CaSR and EV expressing MDA-MB-134 treated with U0124 U0126 and PD98059 for 3 days. Arrows indicate the drug concentration chosen for further experiment (C) Western Blotting using the indicated antibodies on whole cell lysate from CaSR-OE MDA-MB-134 with indicated treatment. (D) Images of crystal violet-stained inserts with CaSR-OE or EV MDA-MB-134 from chemotaxis toward FBS after 72 hours with treatments of MEK inhibitors (U0126 and its negative control U0124 (1 μM) and PD98059 (5 μM)). MEK inhibition prevents CaCl_2_ induced chemotaxis toward FBS in CaSR-OE MDA-MB-134. Data representative of 2 experiments (n=3). Quantification in Figure 5E. (E) Quantification of confocal microscopy imaging shown in Figure 5F of F-actin staining (green) in CaSR-OE and EV MDA-MB-134. CaCl_2_ treatment enhanced F-actin spreading per cell while NPS abrogated the effect. Hoechst staining (blue) was used for nuclear staining. 50µm scale bar. ImageJ software was used to quantify fluorescence intensity per cell. Graphs shows representative data from 5 field per conditions (n=5). Each experiment has been repeated at least 2 times (n=3). P-values are from unpaired t-test analysis. *, P ≤ 0.05. **, P ≤ 0.005. ***, P ≤ 0.001

**Supplementary Figure 10. Estrogen enhances CaSR effect in chemotactic transwell migration**

(A) Enrichment plots show “Estrogen Response Early” (Left panel) and “Estrogen Response Late” (Right panel) gene signature enrichment comparison between CaSR high and low ovarian metastatic tumors. CaSR high and low groups were divided by median expression. (B, C) Images of crystal violet-stained inserts with CaSR-OE and EV MDA-MB-134 with indicated treatment from chemotaxis toward FBS. (B) Inhibition of ER signaling with fulvestrant (ICI), and tamoxifen (4OH) blocks Activated CaSR induced migration by CaCl_2_. (C) In hormone deprived cells, E2 is required for CaSR induced chemotaxis toward FBS with CaCl_2_. (D) Venn diagram shows the intersection of estrogen response genes specifically enriched in MCF7, T47D and MDA-MB-134 cell lines after E2 treatment and enriched in CaSR higher ovarian metastatic tumors. Estrogen response genes were derived from overlapped core enriched genes from gene set enrichment analysis of “Estrogen Response Early” and “Estrogen Response Late” in vehicle vs. E2 groups (cell lines) and CaSR high vs. low groups (ovarian mets).

## Supplementary Data Legends

**Supplementary Data 1.** Cravat Annotated Mutations

**Supplementary Data 2.** PAM50 and Purity Annotations

**Supplementary Data 3.** DEGs ILC vs IDC Ovary Metastasis

**Supplementary Data 4.** DEGs Normal Ovary vs Primary Tumor

**Supplementary Data 5.** DEGs Ovary Metastasis vs Primary Tumor

**Supplementary Data 6.** DEGs Ovary Metastasis vs Primary Tumor

**Supplementary Data 7.** KEGG GSEA DEGs in ILC and IDC

**Supplementary Data 8.** IPA – Ovary Metastasis vs Primary Tumor

## Supplementary Table Legends

**Supplementary Table 1.** Detailed clinical characteristics of the ovarian metastasis cohort.

**Supplementary Table 2.** qPCR Primers

**Supplementary Table 3.** Antibodies

## References

1. Li, C.I., et al., Changing incidence rate of invasive lobular breast carcinoma among older women. Cancer, 2000. 88(11): p. 2561–9.

2. Biglia, N., et al., Increased incidence of lobular breast cancer in women treated with hormone replacement therapy: implications for diagnosis, surgical and medical treatment. Endocr Relat Cancer, 2007. 14(3): p. 549–67.

3. Giaquinto, A.N., et al., *Lobular breast cancer statistics*, *2025*. Cancer, 2025. 131(20): p. e70061.

4. Hilleren, D.J., et al., Invasive lobular carcinoma: mammographic findings in a 10-year experience. Radiology, 1991. 178(1): p. 149–54.

5. Engstrom, M.J., et al., Invasive lobular breast cancer: the prognostic impact of histopathological grade, E-cadherin and molecular subtypes. Histopathology, 2015. 66(3): p. 409–19.

6. Desmedt, C., et al., Genomic Characterization of Primary Invasive Lobular Breast Cancer. J Clin Oncol, 2016. 34(16): p. 1872–81.

7. Adachi, Y., et al., Comparison of clinical outcomes between luminal invasive ductal carcinoma and luminal invasive lobular carcinoma. BMC Cancer, 2016. 16: p. 248.

8. Pestalozzi, B.C., et al., Distinct clinical and prognostic features of infiltrating lobular carcinoma of the breast: combined results of 15 International Breast Cancer Study Group clinical trials. J Clin Oncol, 2008. 26(18): p. 3006–14.

9. Harris, M., et al., A comparison of the metastatic pattern of infiltrating lobular carcinoma and infiltrating duct carcinoma of the breast. Br J Cancer, 1984. 50(1): p. 23–30.

10. Arpino, G., et al., Infiltrating lobular carcinoma of the breast: tumor characteristics and clinical outcome. Breast Cancer Res, 2004. 6(3): p. R149–56.

11. Blohmer, M., et al., Patient treatment and outcome after breast cancer orbital and periorbital metastases: a comprehensive case series including analysis of lobular versus ductal tumor histology. Breast Cancer Res, 2020. 22(1): p. 70.

12. He, H., et al., Distant metastatic disease manifestations in infiltrating lobular carcinoma of the breast. AJR Am J Roentgenol, 2014. 202(5): p. 1140–8.

13. Mathew, A., et al., Distinct Pattern of Metastases in Patients with Invasive Lobular Carcinoma of the Breast. Geburtshilfe Frauenheilkd, 2017. 77(6): p. 660–666.

14. Bigorie, V., et al., Ovarian metastases from breast cancer: report of 29 cases. Cancer, 2010. 116(4): p. 799–804.

15. Lee, S.J., et al., Clinical characteristics of metastatic tumors to the ovaries. J Korean Med Sci, 2009. 24(1): p. 114–9.

16. Pimentel, C., et al., Ovarian Metastases from Breast Cancer: A Series of 28 Cases. Anticancer Res, 2016. 36(8): p. 4195–200.

17. Tian, W., et al., Ovarian metastasis from breast cancer: a comprehensive review. Clin Transl Oncol, 2019. 21(7): p. 819–827.

18. Li, A., et al., Ovarian Metastasis from Invasive Lobular Carcinoma of the Breast: A 6-Case Series with Emphasis on Diagnostic Challenges and the Value of Biopsy. Diagnostics (Basel), 2026. 16(13).

19. Berx, G., et al., E-cadherin is a tumour/invasion suppressor gene mutated in human lobular breast cancers. EMBO J, 1995. 14(24): p. 6107–15.

20. Teo, K., et al., E-cadherin loss induces targetable autocrine activation of growth factor signalling in lobular breast cancer. Sci Rep, 2018. 8(1): p. 15454.

21. Vos, C.B., et al., E-cadherin inactivation in lobular carcinoma in situ of the breast: an early event in tumorigenesis. Br J Cancer, 1997. 76(9): p. 1131–3.

22. Brown, E.M., et al., Cloning and characterization of an extracellular Ca(2+)-sensing receptor from bovine parathyroid. Nature, 1993. 366(6455): p. 575–80.

23. Kim, W. and J.J. Wysolmerski, Calcium-Sensing Receptor in Breast Physiology and Cancer. Front Physiol, 2016. 7: p. 440.

24. Tuffour, A., et al., Role of the calcium-sensing receptor (CaSR) in cancer metastasis to bone: Identifying a potential therapeutic target. Biochim Biophys Acta Rev Cancer, 2021. 1875(2): p. 188528.

25. Sarkar, P. and S. Kumar, Calcium sensing receptor modulation for cancer therapy. Asian Pac J Cancer Prev, 2012. 13(8): p. 3561–8.

26. Savariau, L.A., Jennifer M.; Quin, Ye; Shah, Osama; Basudan, Ahmed Mohammed; McGinn, Olivia; Li, Zheqi; Liu, Tiantong; Tasdemir, Nilgun; Tallapaneni, Pooja; Coffman, Lan; Elishaev, Esther; Lucas, Peter C; Lee, Adrian V; Oesterreich, Steffi, IHC & E-cadherin or Dual staining of paired and orphan primary breast cancer and ovarian metastatic samples. figshare, 10.6084/m9.figshare.17146436, 2022.

27. Levine, K.M., et al., FGFR4 overexpression and hotspot mutations in metastatic ER+ breast cancer are enriched in the lobular subtype. NPJ Breast Cancer, 2019. 5: p. 19.

28. Vareslija, D., et al., Transcriptome Characterization of Matched Primary Breast and Brain Metastatic Tumors to Detect Novel Actionable Targets. J Natl Cancer Inst, 2019. 111(4): p. 388–398.

29. Zhu, L., et al., Metastatic breast cancers have reduced immune cell recruitment but harbor increased macrophages relative to their matched primary tumors. J Immunother Cancer, 2019. 7(1): p. 265.

30. Priedigkeit, N., et al., Exome-capture RNA sequencing of decade-old breast cancers and matched decalcified bone metastases. JCI Insight, 2017. 2(17).

31. Smith, N.G., et al., Targeted mutation detection in breast cancer using MammaSeq. Breast Cancer Res, 2019. 21(1): p. 22.

32. Chakravarty, D., et al., OncoKB: A Precision Oncology Knowledge Base. JCO Precis Oncol, 2017. 2017.

33. Huang, L., et al., The cancer precision medicine knowledge base for structured clinical-grade mutations and interpretations. J Am Med Inform Assoc, 2017. 24(3): p. 513–519.

34. Basudan, A., et al., Frequent ESR1 and CDK Pathway Copy-Number Alterations in Metastatic Breast Cancer. Molecular Cancer Research, 2019. 17(2): p. 457–468.

35. Cejalvo, J.M., et al., Intrinsic Subtypes and Gene Expression Profiles in Primary and Metastatic Breast Cancer. Cancer Res, 2017. 77(9): p. 2213–2221.

36. Mellouli, M., et al., Discordance in receptor status between primary and metastatic breast cancer and overall survival: A single-center analysis. Ann Diagn Pathol, 2022. 61: p. 152044.

37. Garcia-Recio, S., et al., FGFR4 regulates tumor subtype differentiation in luminal breast cancer and metastatic disease. J Clin Invest, 2020. 130(9): p. 4871–4887.

38. Klebe, M., et al., Frequent Molecular Subtype Switching and Gene Expression Alterations in Lung and Pleural Metastasis From Luminal A-Type Breast Cancer. JCO Precis Oncol, 2020. 4.

39. Langmia, I.M., et al., CYP2B6 Functional Variability in Drug Metabolism and Exposure Across Populations-Implication for Drug Safety, Dosing, and Individualized Therapy. Front Genet, 2021. 12: p. 692234.

40. Zeng, Q., et al., Synaptic proximity enables NMDAR signalling to promote brain metastasis. Nature, 2019. 573(7775): p. 526–531.

41. Stires, H., et al., Integrated molecular analysis of Tamoxifen-resistant invasive lobular breast cancer cells identifies MAPK and GRM/mGluR signaling as therapeutic vulnerabilities. Mol Cell Endocrinol, 2018. 471: p. 105–117.

42. Young, T.A., et al., Glutamate Transport Proteins and Metabolic Enzymes are Poor Prognostic Factors in Invasive Lobular Carcinoma. bioRxiv, 2024.

43. Brown, E.M., et al., Cloning and characterization of an extracellular Ca(2+)-sensing receptor from parathyroid and kidney: new insights into the physiology and pathophysiology of calcium metabolism. Nephrol Dial Transplant, 1994. 9(12): p. 1703–6.

44. Conigrave, A.D. and D.T. Ward, Calcium-sensing receptor (CaSR): pharmacological properties and signaling pathways. Best Pract Res Clin Endocrinol Metab, 2013. 27(3): p. 315–31.

45. Das, S., et al., The CaSR in Pathogenesis of Breast Cancer: A New Target for Early Stage Bone Metastases. Front Oncol, 2020. 10: p. 69.

46. Yang, Y. and B. Wang, PTH1R-CaSR Cross Talk: New Treatment Options for Breast Cancer Osteolytic Bone Metastases. Int J Endocrinol, 2018. 2018: p. 7120979.

47. Xie, W., H. Xu, and Y. Cheng, Calcium-sensing Receptor, a Potential Biomarker Revealed by Large-scale Public Databases and Experimental Validation in Breast Cancer. Technol Cancer Res Treat, 2024.

48. Hannan, F.M., et al., The calcium-sensing receptor in physiology and in calcitropic and noncalcitropic diseases. Nat Rev Endocrinol, 2018. 15(1): p. 33–51.

49. Saidak, Z., et al., Extracellular calcium promotes the migration of breast cancer cells through the activation of the calcium sensing receptor. Exp Cell Res, 2009. 315(12): p. 2072–80.

50. Tasdemir, N., et al., Comprehensive Phenotypic Characterization of Human Invasive Lobular Carcinoma Cell Lines in 2D and 3D Cultures. Cancer Res, 2018. 78(21): p. 6209–6222.

51. Davies, S.L., et al., Ca2+-sensing receptor induces Rho kinase-mediated actin stress fiber assembly and altered cell morphology, but not in response to aromatic amino acids. Am J Physiol Cell Physiol, 2006. 290(6): p. C1543–51.

52. Webb, D.J., J.T. Parsons, and A.F. Horwitz, Adhesion assembly, disassembly and turnover in migrating cells -- over and over and over again. Nat Cell Biol, 2002. 4(4): p. E97–100.

53. Tanimura, S. and K. Takeda, ERK signalling as a regulator of cell motility. J Biochem, 2017. 162(3): p. 145–154.

54. Dudley, D.T., et al., A synthetic inhibitor of the mitogen-activated protein kinase cascade. Proc Natl Acad Sci U S A, 1995. 92(17): p. 7686–9.

55. Favata, M.F., et al., Identification of a novel inhibitor of mitogen-activated protein kinase kinase. J Biol Chem, 1998. 273(29): p. 18623–32.

56. Journe, F., et al., Extracellular calcium downregulates estrogen receptor alpha and increases its transcriptional activity through calcium-sensing receptor in breast cancer cells. Bone, 2004. 35(2): p. 479–88.

57. Chaffer, C.L. and R.A. Weinberg, A perspective on cancer cell metastasis. Science, 2011. 331(6024): p. 1559–64.

58. Kutasovic, J.R., et al., Breast cancer metastasis to gynaecological organs: a clinico-pathological and molecular profiling study. J Pathol Clin Res, 2019. 5(1): p. 25–39.

59. Cejalvo, J.M., et al., Clinical implications of routine genomic mutation sequencing in PIK3CA/AKT1 and KRAS/NRAS/BRAF in metastatic breast cancer. Breast Cancer Res Treat, 2016. 160(1): p. 69–77.

60. Razavi, P., et al., The Genomic Landscape of Endocrine-Resistant Advanced Breast Cancers. Cancer Cell, 2018. 34(3): p. 427–438 e6.

61. Yates, L.R., et al., Genomic Evolution of Breast Cancer Metastasis and Relapse. Cancer Cell, 2017. 32(2): p. 169–184 e7.

62. Ma, J., et al., Targeting of erbB3 receptor to overcome resistance in cancer treatment. Mol Cancer, 2014. 13: p. 105.

63. Sutherland, R.L., Endocrine resistance in breast cancer: new roles for ErbB3 and ErbB4. Breast Cancer Research, 2011. 13(3): p. 106.

64. Tomlinson, D.C., M.A. Knowles, and V. Speirs, Mechanisms of FGFR3 actions in endocrine resistant breast cancer. Int J Cancer, 2012. 130(12): p. 2857–66.

65. Agashe, R.P., S.M. Lippman, and R. Kurzrock, JAK: Not Just Another Kinase. Mol Cancer Ther, 2022. 21(12): p. 1757–1764.

66. Raivola, J., et al., Hyperactivation of Oncogenic JAK3 Mutants Depend on ATP Binding to the Pseudokinase Domain. Front Oncol, 2018. 8: p. 560.

67. Zhou, R., et al., Clinical Impact of 11q13.3 Amplification on Immune Cell Infiltration and Prognosis in Breast Cancer. Int J Gen Med, 2022. 15: p. 4037–4052.

68. Christgen, M., et al., Inter-observer agreement for the histological diagnosis of invasive lobular breast carcinoma. J Pathol Clin Res, 2022. 8(2): p. 191–205.

69. Mehta, M.S., et al., Metabotropic glutamate receptor 1 expression and its polymorphic variants associate with breast cancer phenotypes. PLoS One, 2013. 8(7): p. e69851.

70. Ribeiro, M.P., J.B. Custodio, and A.E. Santos, Ionotropic glutamate receptor antagonists and cancer therapy: time to think out of the box? Cancer Chemother Pharmacol, 2017. 79(2): p. 219–225.

71. Brennan, S.C. and A.D. Conigrave, Regulation of cellular signal transduction pathways by the extracellular calcium-sensing receptor. Curr Pharm Biotechnol, 2009. 10(3): p. 270–81.

72. Zavala-Barrera, C., et al., The calcium sensing receptor (CaSR) promotes Rab27B expression and activity to control secretion in breast cancer cells. Biochim Biophys Acta Mol Cell Res, 2021. 1868(7): p. 119026.

73. Bellet, M., et al., Twelve-Month Estrogen Levels in Premenopausal Women With Hormone Receptor-Positive Breast Cancer Receiving Adjuvant Triptorelin Plus Exemestane or Tamoxifen in the Suppression of Ovarian Function Trial (SOFT): The SOFT-EST Substudy. J Clin Oncol, 2016. 34(14): p. 1584–93.

74. Hindson, B.J., et al., High-throughput droplet digital PCR system for absolute quantitation of DNA copy number. Anal Chem, 2011. 83(22): p. 8604–10.

75. Liang, C.C., A.Y. Park, and J.L. Guan, In vitro scratch assay: a convenient and inexpensive method for analysis of cell migration in vitro. Nat Protoc, 2007. 2(2): p. 329–33.

76. Sikora, M.J., et al., Mechanisms of estrogen-independent breast cancer growth driven by low estrogen concentrations are unique versus complete estrogen deprivation. Breast Cancer Res Treat, 2012. 134(3): p. 1027–39.

77. Bahreini, A., et al., Mutation site and context dependent effects of ESR1 mutation in genome-edited breast cancer cell models. Breast Cancer Res, 2017. 19(1): p. 60.

78. Sikora, M.J., et al., Invasive lobular carcinoma cell lines are characterized by unique estrogen- mediated gene expression patterns and altered tamoxifen response. Cancer Res, 2014. 74(5): p. 1463–74.

