## Supplementary material for "Breast cancer ovarian metastases show increased activity of GPCR pathways": Suppl figures

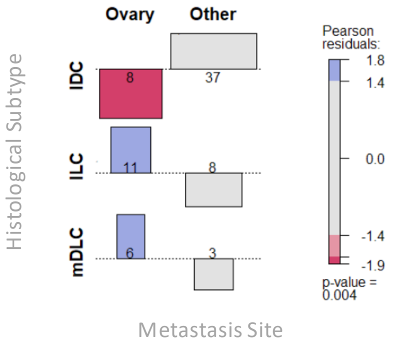


**Supplementary Figure 1. Association between ovarian metastasis and histological subtype**

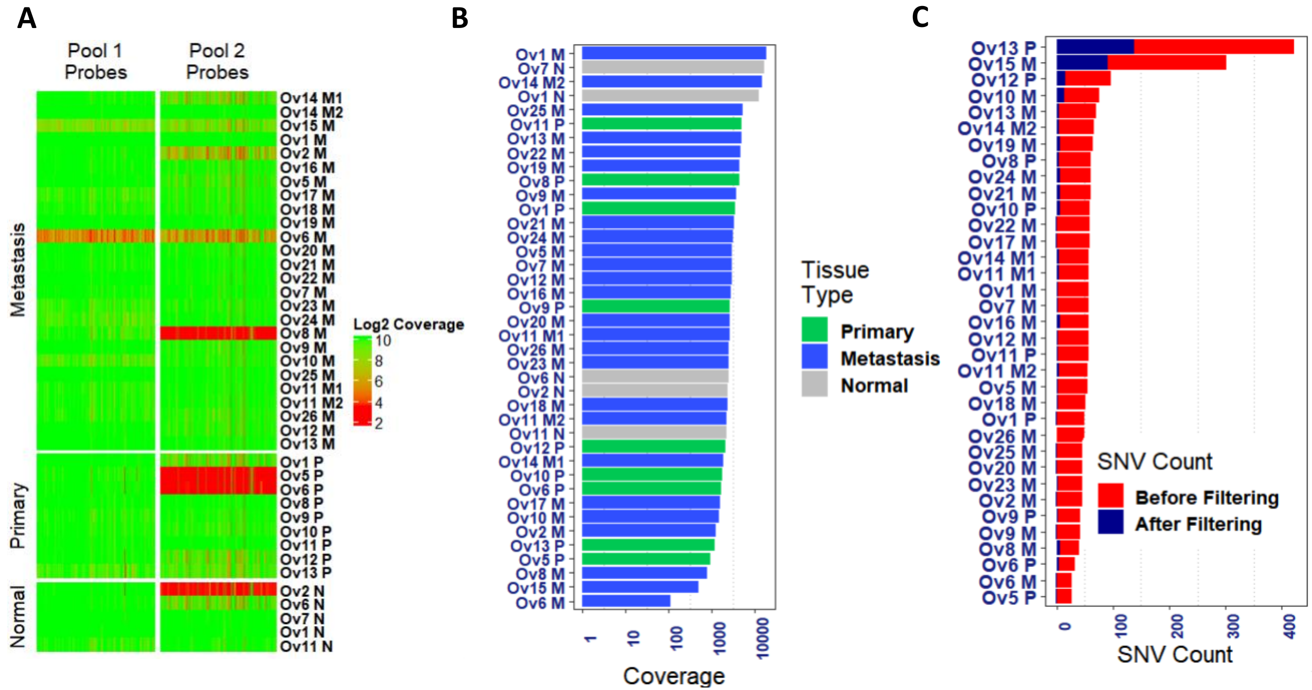


**Supplementary Figure 2: MammaSeq™ based DNAseq QC**

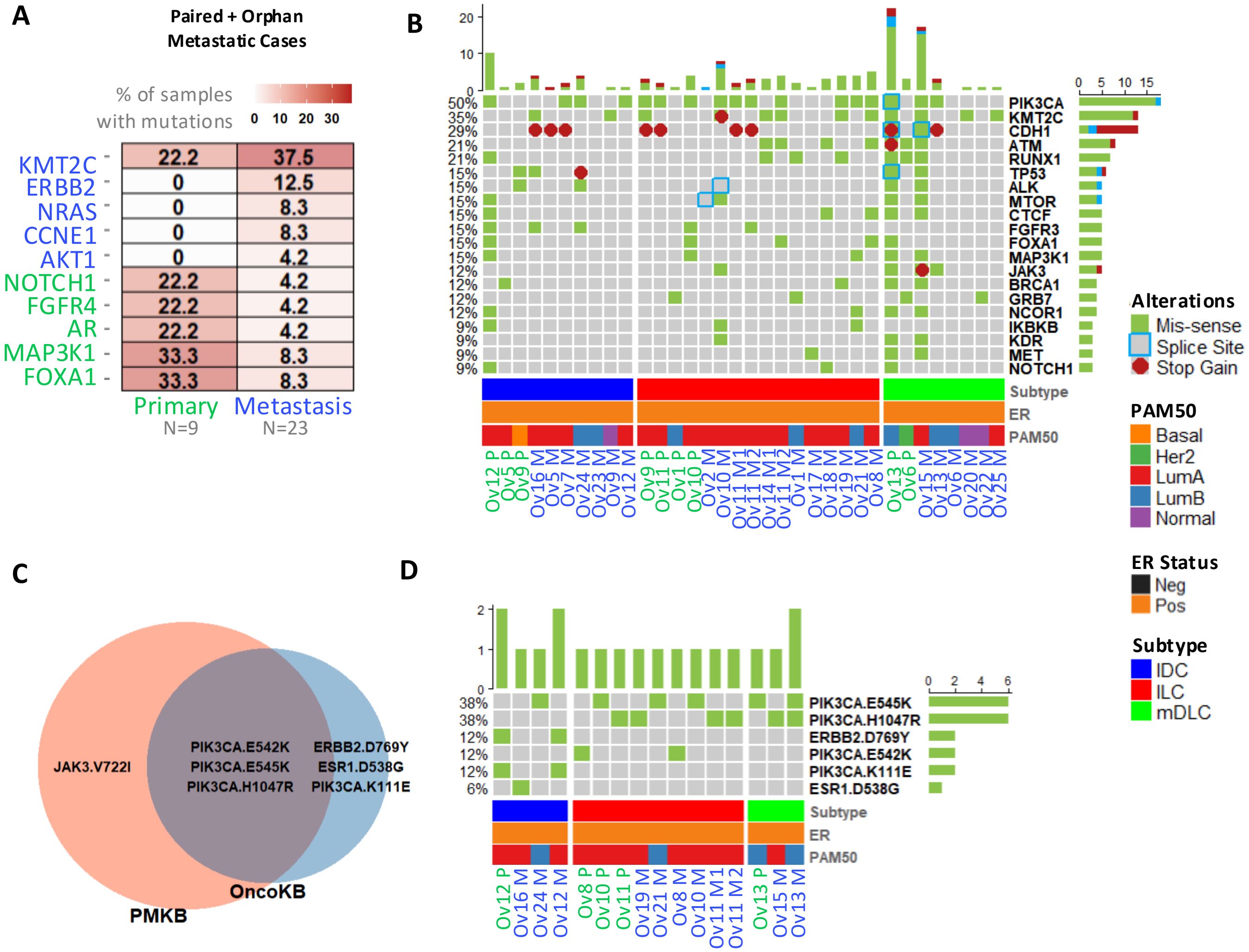


**Supplementary Figure 3. Mutational landscape of all (paired + orphan) ovarian metastatic samples and clinical actionability analysis**

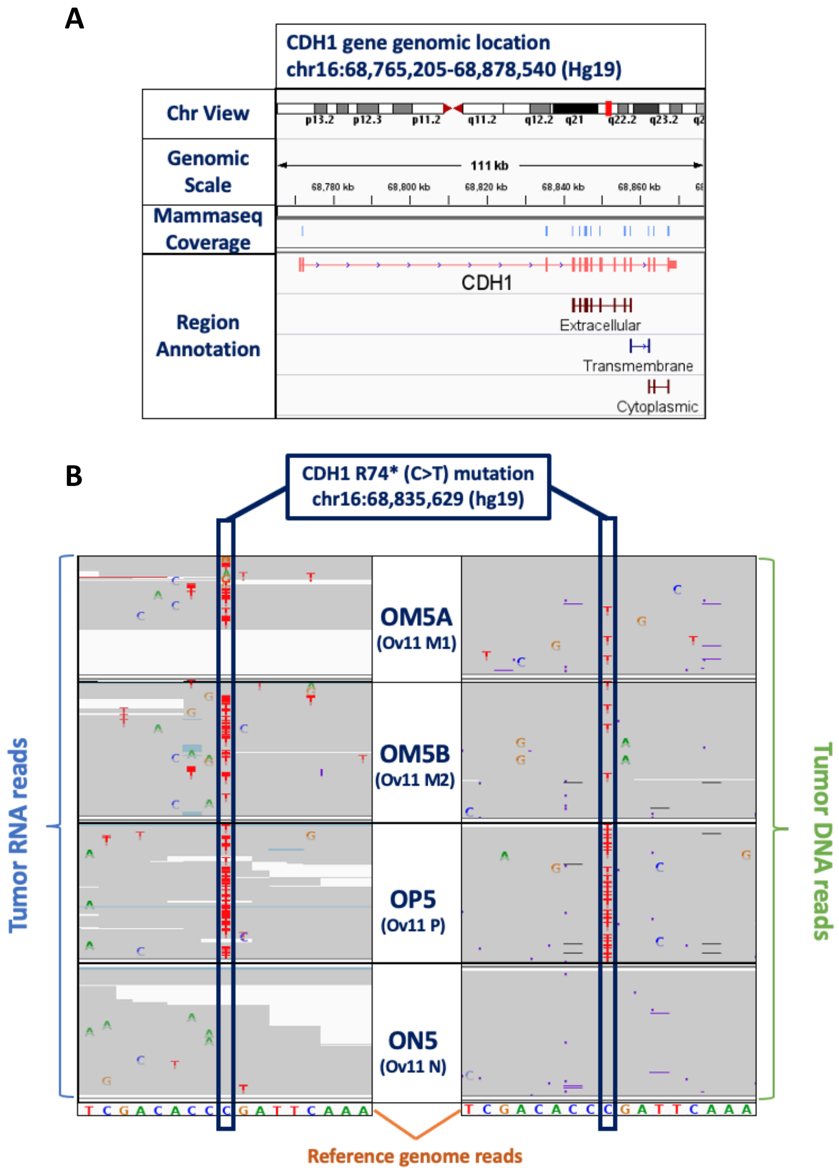


**Supplementary Figure 4. Ov11 CDH1 R74* mutation assessment in RNA/DNA reads**

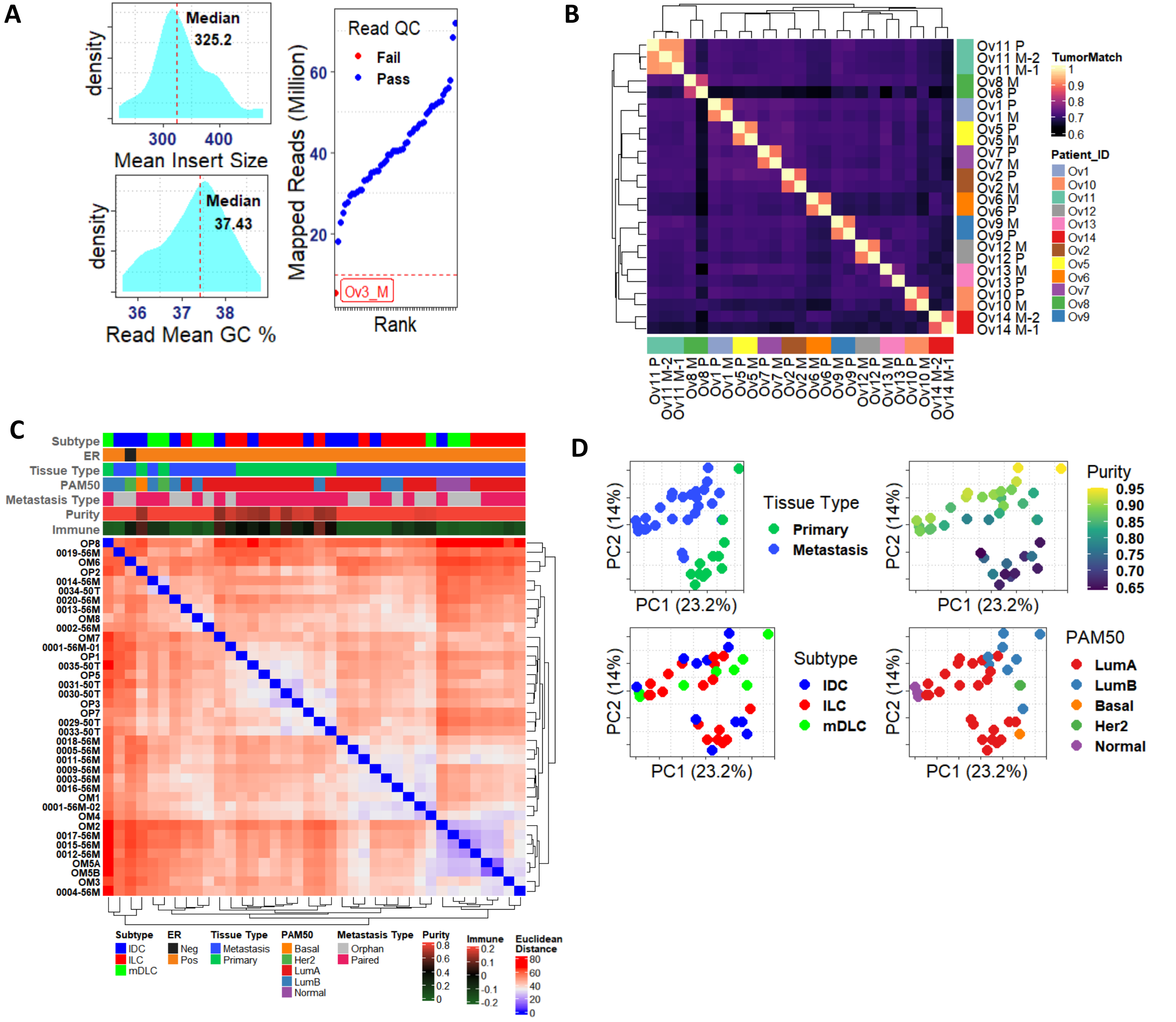


**Supplementary Figure 5. RNAseq QC and exploratory analysis**

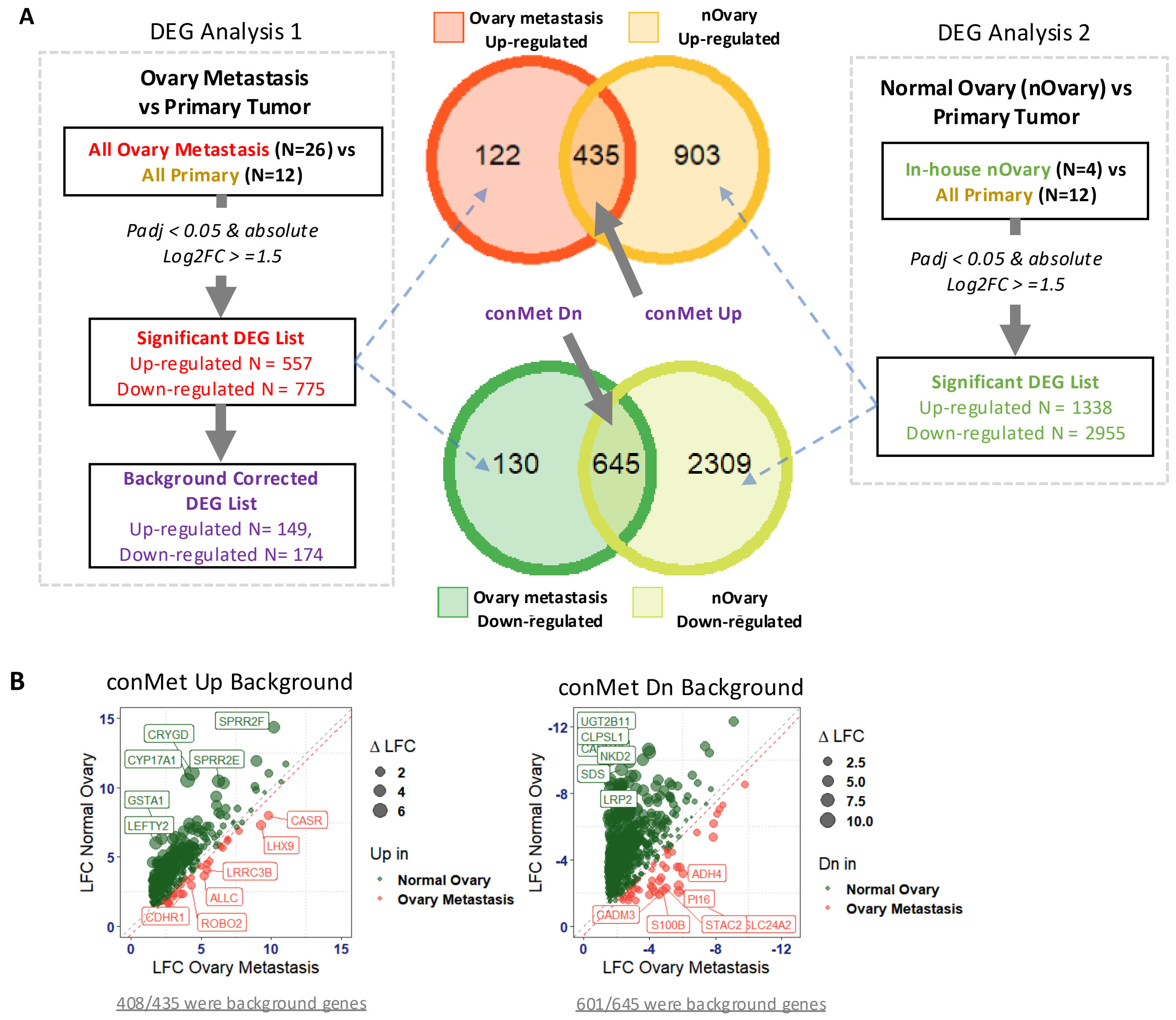


**Supplementary Figure 6. DEA scheme for identification of ovarian metastasis associated DEG**

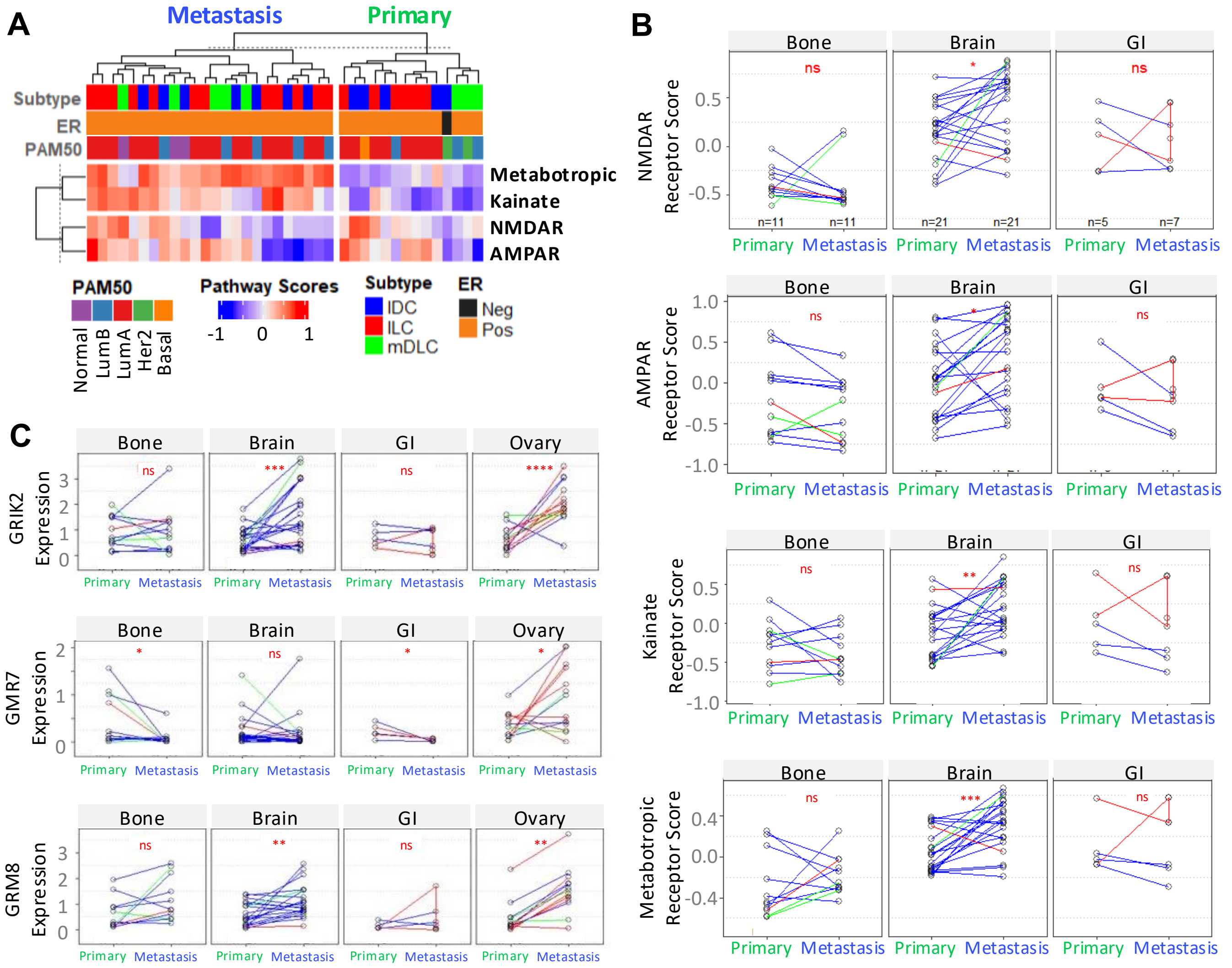


**Supplementary Figure 7. Glutamate receptor family signaling activity in ovary, bone, GI and brain metastasis**

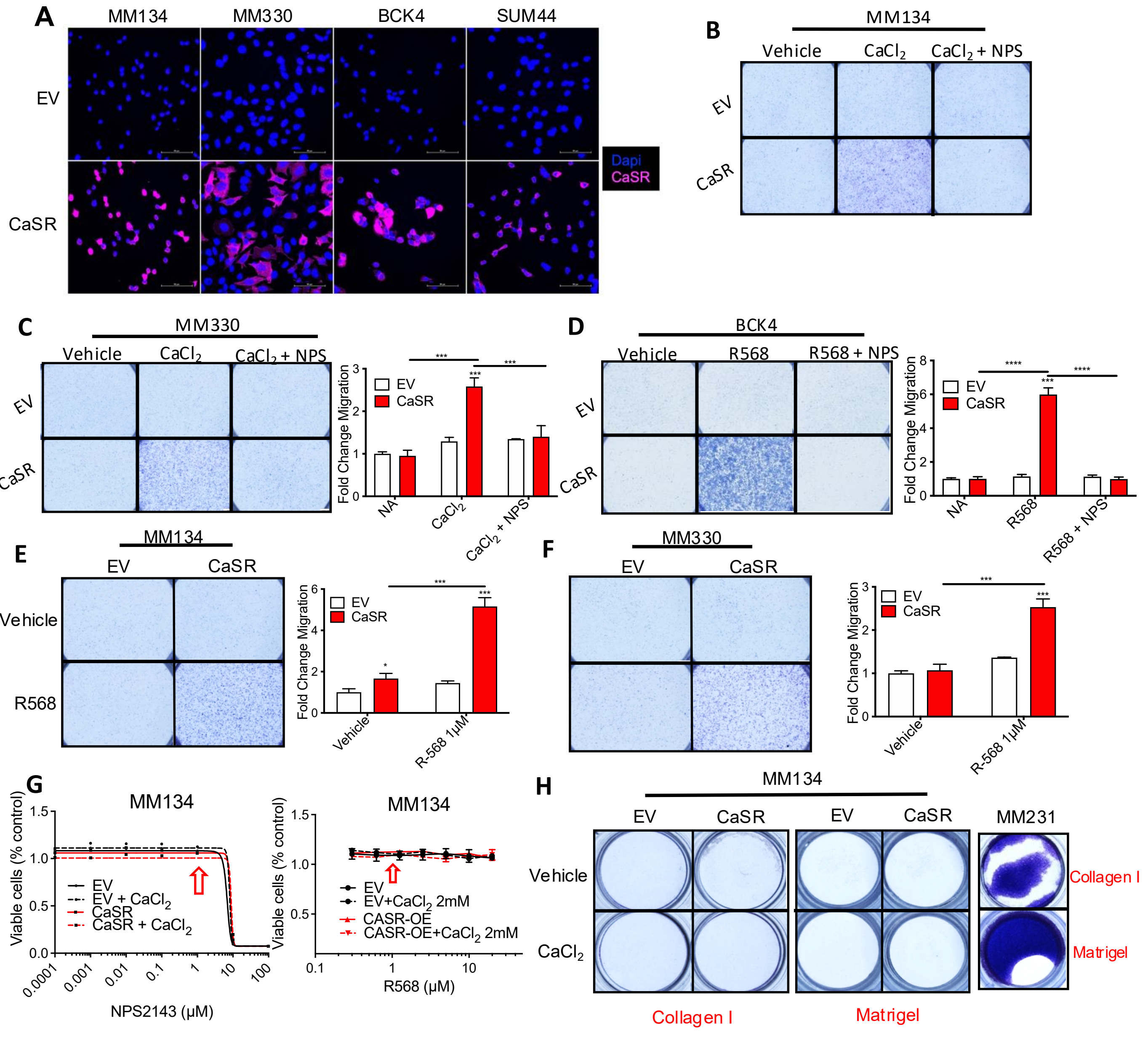


**Supplementary Figure 8. CaSR overexpression and calcimimetic activation promote migration and growth in ILC cell lines, reversed by calcilytic inhibition**

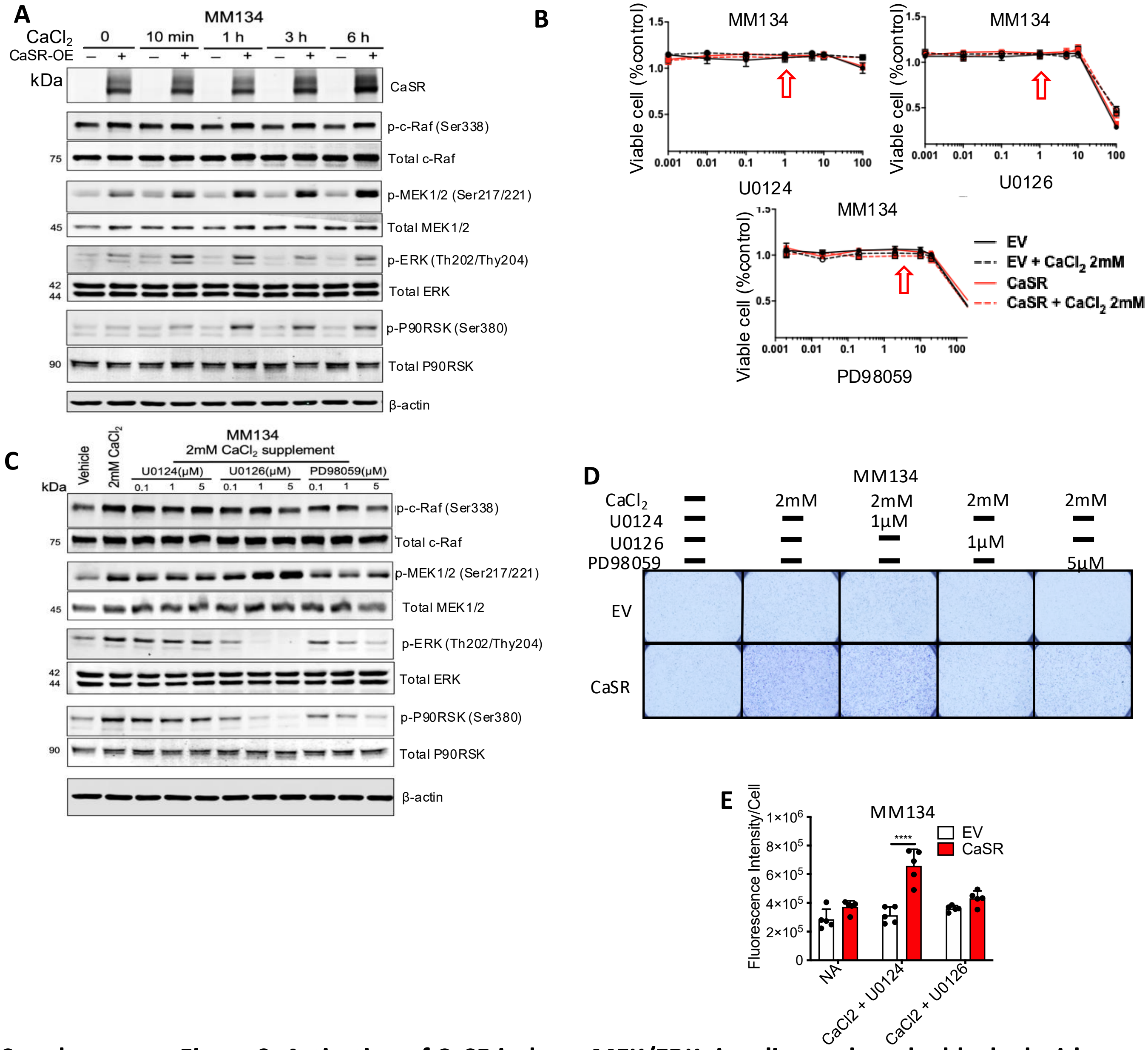


**Supplementary Figure 9. Activation of CaSR induces MEK/ERK signaling and can be blocked with calcilytic**

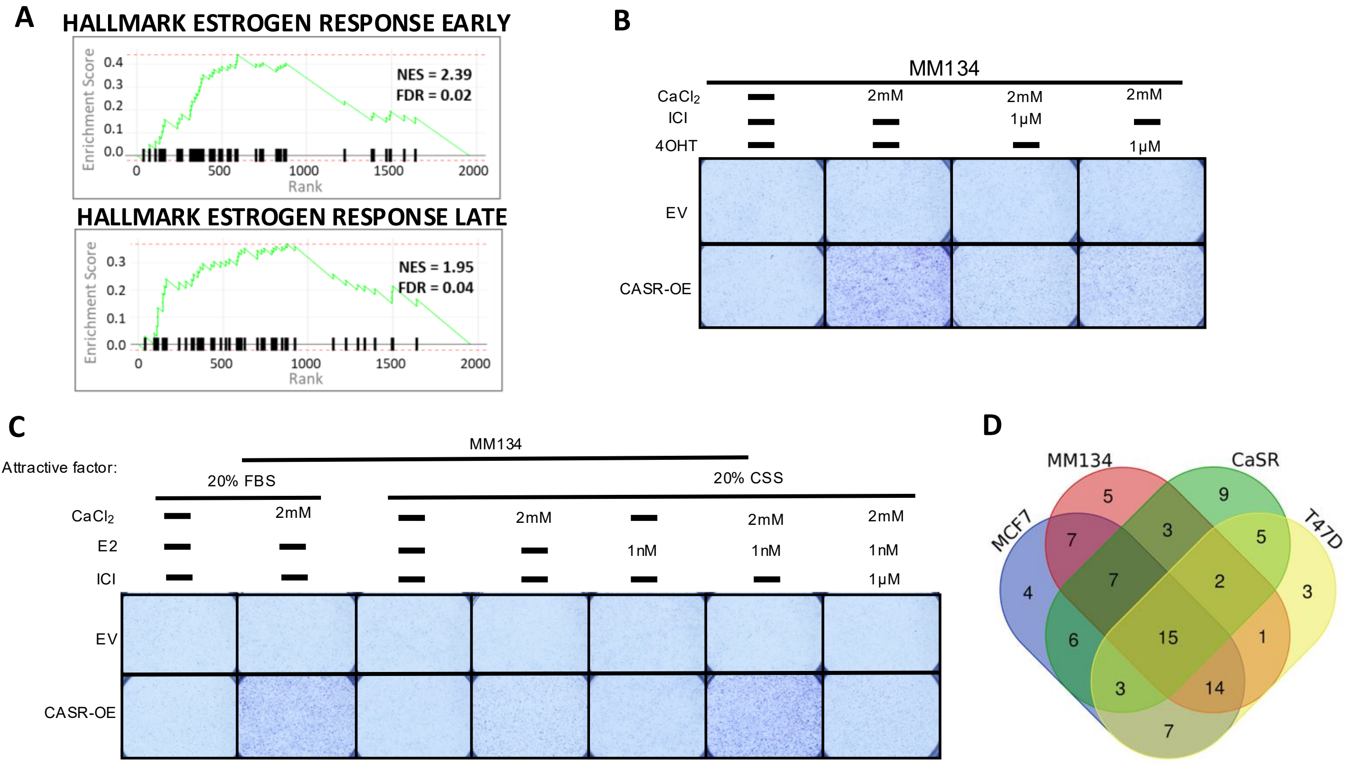


**Supplementary Figure 10. Estrogen enhance CaSR effect in chemotactic transwell migration**
