## Supplementary material for "Breast cancer ovarian metastases show increased activity of GPCR pathways": Suppl Methods

#### Supplementary Methods

### Supplementary Methods

#### ***RNA sequencing***

##### *1. Exome-capture library preparation and RNA sequencing*

Library preparation using TruSeq RNA Access library preparation (Illumina), and sequencing was performed at the Genomic Core Facility at Children's Hospital of UPMC. RNA samples were sonication-fragmented, followed by cDNA synthesis, adapter ligation, and PCR amplification. Lastly, biotinylated probes were hybridized to the target regions (exome), followed by capture step using streptavidin beads, then elution of the beads. NextSeq500 platform was used to produce 2 x 75bp paired end reads with target reads of 40-50 million.

##### *2. RNA-sequencing expression quantification and normalization*

RNA expression quantification and gene-level summarization were performed using Salmon v0.12.0 [1] and tximport v1.16.1[2], respectively. Briefly, Salmon quasi-mapping was performed using 31-kmer transcriptome index generated using GRCh38 Ensemble v82 transcript annotation. Resulting expression estimates were summarized to gene-level by tximport and used in subsequent analyses. Moreover, RNAseq read alignments were also generated using STAR v2.4.2a [3] for quality assessment as well as variant calling as described below.

##### *3. RNA-seq quality assessment*

Basic quality control (QC) metrics like read mapping were assessed using Salmon met_info file output. QoRTs v1.1.8 [4] was used to generate a detailed QC report using STAR generated BAM files as input. QoRTs QC data was visualized using ggplot2 [5] package in R. One RNAseq sample (0006-56M) was removed from analysis due to small number of reads (< 10 million reads).

##### *4. Variant calling and genotype matching*

RNAseq variants were identified using GATK 3 pipeline as per their best practices guidelines [6]. Resulting vcf files were used to test kinship (genotype relation) between samples to confirm patient-matched samples were indeed from same patient using our previously published custom R script tool “tumorMatch” [7]. Briefly, the genotype calls for all variants were compared to compute proportion of shared variants (POSV) across all samples. The POSV scores were visualized as heatmap using ComplexHeatmap [8].

##### *5. Intrinsic PAM50 molecular subtyping analysis*

To make accurate PAM50 calls it is important that the classification cohort’s ER status distribution resembles that of the training cohort used to define PAM50 centroids[9]. For ovary metastasis cohort, ER status was defined using ER IHC staining. ER status was further refined using ESR1 mRNA levels with respect to ESR1 cut off for ER+ and ER- samples in breastMETs cohort, respectively. Three ovary metastasis cohort samples; Ov13 M (ER IHC N/A), Ov12 M (ER IHC N/A) and Ov12 P (IHC ER-) were reclassified to ER+, ER- and ER+, respectively (see Supplementary data 2 –PAM50 and tumor purity annotation columns ER IHC and ER inferred from mRNA). PAM50 subtyping was performed using genefu [9] as previously described [7]. Briefly, test samples were selected randomly such that even distribution of ER+ and ER- tumors was present. This process was repeated to get 20 ER-balanced sample subsets. PAM50 gene expressions for each subset were provided as input to *molecular.subtyping* function in genefu. This resulted in 20 PAM50 calls and call probability scores per sample. For each sample, the final PAM50 assignment was defined as mode of 20 calls while the final PAM50 probability score was an average of all 20 call probability scores. Subtype switching was defined as at least one of the PAM50 subtype call probability scores between paired primary and metastatic samples differed by 0.25 or 25%. Associations between metastatic sites and primary tumor histology were tested using Pearson's chi square test with vcd package [10].

##### 6. ***Purity analysis***

Tumor purity analysis was performed using Estimate [11, 12]. Immune scores were computed using consensusTME package in R [11].

##### 7. ***Additional RNAseq datasets***

The Genotype-Tissue Expression Project (GTEx) [13] normal tissue RNAseq counts were downloaded from gtexportal (https://gtexportal.org/home/datasets). The paired primary breast tumor to brain, bone, and GI metastasis (breastMETs) RNAseq counts used in this study were acquired from our previous publications [7, 14, 15].

##### 8. ***Differential Expression Analysis (DEA)***

DEA between ILC (N=12) vs IDC (N=8) ovarian metastasis was performed using following design: ~Subtype [i.e., ILC or IDC]. Significant DEGs were defined as those with absolute log2 fold change >= 1.5 and FDR-adjusted P value of < 0.05 (DEGs are listed in supplementary data 3).

Two DEA were performed to identify background corrected ovarian metastasis DEGs; DEA 1 to identify DEGs associated with ovary metastasis (Supplementary Data 6) and DEA 2 to identify DEGs associated with normal ovary i.e., background (Supplementary Data 7, and Supplementary Figure. S6A). Given our cohort comprised of paired metastatic samples as well, a multifactor design was implemented for DEA 1 as following ~TissueType [i.e., Metastasis or Primary Tumor] + Patient_ID. For DEA 2 following design was implemented ~TissueType [i.e., Normal Ovary or Primary Tumor]. Significant DEGs were defined as described above. Ovary metastasis DEGs potentially contaminated by background were identified as such; contamination of metastasis up (conMet Up) geneset (N = 435) was defined as overlap of up-regulated DEGs in both ovary metastasis (from DEA 1) and normal ovary (from DEA 2). Similarly, contamination of metastasis down (conMet Dn) geneset (N = 645) was defined as overlap of downregulated genes in both ovary metastasis (from DEA 1) and normal ovary (from DEA 2). Both conMet Up and Dn genesets underwent background correction by removing genes with 0.5 higher and lower log2FC difference, respectively, between DEA 1 and DEA 2 (Supplementary Figure. S6B, C, Supplementary Data 8). DEGs were visualized using volcano plots and heatmaps were generated using ggplot2 [5] and ComplexHeatmap [8] packages in R, respectively. Venn diagrams showing overlap between DGE analysis 1 and 2 were generated using seqsetvis package in R [16]. All DEA analysis were performed using DESeq2 package in R [17] and are listed in the Supplementary Data 3, 6, 7, 8.

##### 9. ***Clustering***

Clustering in sample-sample similarity heatmaps and DEG heatmaps was performed using ComplexHeatmap [8] using default parameters (Euclidean distance and complete linkage).

##### *10.* ***Pathway analysis***

Gene set enrichment analysis (GSEA) of DEGs with higher expression in IDC and ILC metastasis using KEGG pathways was performed using HypeR package in R [18].

GSEA of “Estrogen Response Early” and “Estrogen Response Late” Hallmark signature were performed using GSEA 4.0.3 software [19]. For this test, 25 ER+ ovary metastatic cases were divided into high and low groups by the median of log2(TPM+1) values of CASR expression.

Ingenuity Pathway Analysis (QIAGEN Inc., https://www.qiagenbioinformatics.com/products/ingenuity-pathway-analysis) was performed on corrected ovarian metastasis vs primary tumor DEG list. Pathway enrichment results were plotted as barplots using R.

Activity of various glutamate receptor family genesets (obtained from Zeng et al study [20]) was scored across samples using gene set variation analysis (GSVA) package in R [21].

***Mutation Analysis***

***1. MammaSeq library preparation and Ion Torrent Sequencing***

Samples with adequate DNA underwent Ion Torrent Sequencing at the University of Pittsburgh Genomics Core as previously described[22, 23]. Briefly, 20ng of DNA was used per sample for library preparation using Ion AmpliSeq library kit (ThermoFisher Scientific) and MammaSeq primer panel. Emulsion polymerase chain reaction (PCR) was used for target enrichment using Ion OneTouch 2 system (ThermoFisher Scientific). The enriched library sequenced using Ion Torrent Personal Genome Machine (PGM, ThermoFisher Scientific).

***2. Preprocessing and* Quality Assessment**

Resulting raw reads were aligned to reference genome Hg19 and underwent variant calling using Ion Torrent Suite V4.0 proprietary pipeline. Captured region and mean sample coverages were analyzed using TarSeqQC v1.18.0 [8, 24]. Sequencing coverage filter was defined as average coverage greater than 100X.

##### ***3. Variant Analysis***

Variant analysis was performed as described previously [22, 23]. Briefly, variant call format (VCF) V4.0 files from Torrent Suite were merged and annotated using CRAVAT [25]. Variant filtering was performed to remove pipeline artifacts, common polymorphisms and enrich for somatic variants. Filtered variants were annotated for clinical significance using PMKB and OncoKB databases [22, 23, 26, 27]. Mutational frequency change analysis was performed by computing mutational frequency of each gene in following groups individually: all primary (mutFreqPri), paired metastatic (mutFreqPairedMet) and all metastatic samples (mutFreqAllMet). Mutational frequency change was defined as mutFreqPairedMet (or mutFreqAllMet) – mutFreqPri. Top 5 enriched (+ive change) and depleted (-ive change) genes were identified based on the mutational frequency change and plotted using ggplot tiles for pairedMet vs primary and allMet vs primary.

##### ***4. Validation of CDH1 R74* mutation***

Supporting reads for CDH1 R74* (C > T) mutation were visualized IGV [28] using BAM files from both RNAseq (generated using STAR as described above) and MammaSeq assay for paired samples; ON5 (normal), OP5 (primary tumor), OM5A (ovary metastasis) and OM5B (ovary metastasis). Validation using ddPCR is described below. Analysis code used to perform bioinformatics analysis is currently being deposited at github (https://github.com/osamashiraz/LO_Lab_CASR_2021) and will be made available at time of publication.

***Nanostring Copy Number (CN) Analysis***

CN data was obtained from our previous study [29] where we characterized CN alterations (focusing on *ESR1*) in n=108 metastases using Nanostring based approach. This study included 23 ovarian metastasis and 6 primary tumor samples overlapping with our cohort. CN data normalized to invariant reference probes was used in our study. CN categories were defined as following amplifications (AMP) >= 5 CN, gains (GAIN) >= 3 CN, loss of heterozygosity (LOH) <= 1.5 CN, deletions (DEL) <= 1 CN.

***Cells and reagents***

Breast cancer cell lines MDA-MB-134-VI (MDA-MB-134), MDA-MB-330 and MDA-MB-231 were obtained from the ATCC. SUM44PE (SUM44) was purchased from Asterand, and we obtained BCK4 cells from Dr. Jacobsen [30]. MDA-MB-134 and MDA-MB-330 cells were maintained in 1:1 DMEM: L-15 media (Life Technology) with 10% FBS. MDA-MB-231 was maintained in DMEM with 10% FBS. SUM44 was maintained as described previously in DMEM-F12 with 2% charcoal-stripped serum and supplements[31]. BCK4 was cultured in MEM media with 10% FBS supplemented by 5ml Non-essential Amino Acid (Life Technologies) and $10^{-9}$ M Insulin (Sigma-Aldrich). ICI182780 (fulvestrant) was purchased from Tocris Bioscience. 17β-Estradiol (E2), 4-hydroxytamoxifen (4-OHT), NPS2143, MEK inhibitors (U0124, U0126, PD-98059) were obtained from Sigma-Aldrich (St. Louis, MO).

***RNA extraction and qPCR***

Total RNA was extracted using RNeasy kit (Qiagen). cDNA conversion used PrimeScritpt RT Master Mix kit (Takara Bio), and quantitative PCR (qPCR) reactions used SsoAdvanced Universal SYBR green (Bio-Rad) on a CFX384 thermocycler (Bio-Rad), according to manufacturer’s instructions. Primer sequences are available in **Supplementary Table S2**.

***Immunoblotting***

Western blotting was performed as previously described[32]. Antibodies used in this study are shown in **Supplementary Table S3**. Nitrocellulose membranes were scanned using the Odyssey Infrared Imaging System (Li-Cor Biosciences, Lincoln, NE).

***Proliferation, migration, transwell chemotaxis and haptotaxis, and invasion assays***

Cellular proliferation assays were performed with PrestoBlue (Invitrogen) according to the manufacturer’s instructions. Wound-scratch assays were performed as described previously using the IncuCyte Zoom Live Cell Imaging System (Essen Bioscience)[33, 34]. The Essen Bioscience Imagelock 96-well plate was coated with 100ug/ml matrigel and left in 37 ºC incubator for 1 to 2 hours prior seeding $5\times10^5$ cells per well. 24 hours later, the wound scratch was made by the wound maker and the plate was placed into the IncuCyte machine.

For transwell experiments, cells were serum starved overnight, then 3 million cells were plated per 8 µm pore size inserts in 500 ul FBS free media and supplements as indicated for each experiment (Fisher Scientific #353097 for chemotaxis toward FBS, EMD Millipore #ECM582 for haptotaxis to Collagen I, Millipore Sigma #ECM551 for ECM invasion). The bottom chamber was filled with 1ml of media with 20% FBS and other supplements as indicated for each experiment. After 72 hours, the inserts were placed onto a fresh plate containing 400ul of 0.5% crystal violet in 40% methanol for 15 min at room temperature. Next, the inserts were thoroughly washed with water, while avoiding the outside of the membrane. The inserts were left to dry and photographed with the Olympus SZX16 dissecting microscope. The crystal violet was dissolved in 150ul 0.1 M sodium citrate solution, which 100ul was used to read the OD560 in the name machine.

We did notice major growth inhibition of BCK4 cells at low $CaCl_2$ concentration suggesting toxicity and we therefore used the calcimimetic R568 for studies in this cell line.

***Hormone Deprivation***

For transwell experiment with CSS treatment cells were hormone-deprived using charcoal-stripped FBS (CSS) (12676, Life Technologies), as described previously [76] in phenol red-free improved minimum essential medium (IMEM) + 10 % CSS (2 % CSS for SUM44PE only).

#### References:


1. Patro, R., et al., *Salmon provides fast and bias-aware quantification of transcript expression.* Nat Methods, 2017. **14**(4): p. 417-419.
2. Soneson, C., M.I. Love, and M.D. Robinson, *Differential analyses for RNA-seq: transcript-level estimates improve gene-level inferences.* F1000Res, 2015. **4**: p. 1521.
3. Dobin, A., et al., *STAR: ultrafast universal RNA-seq aligner.* Bioinformatics, 2013. **29**(1): p. 15-21.
4. Hartley, S.W. and J.C. Mullikin, *QoRTs: a comprehensive toolset for quality control and data processing of RNA-Seq experiments.* BMC Bioinformatics, 2015. **16**: p. 224.
5. H, W., *ggplot2: Elegant Graphics for Data Analysis.* 2016: Springer-Verlag New York.
6. Van der Auwera, G.A., et al., *From FastQ data to high confidence variant calls: the Genome Analysis Toolkit best practices pipeline.* Curr Protoc Bioinformatics, 2013. **43**: p. 11 10 1-11 10 33.
7. Vareslija, D., et al., *Transcriptome Characterization of Matched Primary Breast and Brain Metastatic Tumors to Detect Novel Actionable Targets.* J Natl Cancer Inst, 2019. **111**(4): p. 388-398.
8. Gu, Z., R. Eils, and M. Schlesner, *Complex heatmaps reveal patterns and correlations in multidimensional genomic data.* Bioinformatics, 2016. **32**(18): p. 2847-9.
9. Gendoo, D.M., et al., *Genefu: an R/Bioconductor package for computation of gene expression-based signatures in breast cancer.* Bioinformatics, 2016. **32**(7): p. 1097-9.
10. Zeileis, A.M., D.; Hornik, K., *Residual–Based Shadings for Visualizing (Conditional) Independence.* Journal of Computational and Graphical Statistics, 2007. **Volume 16**.
11. Jimenez-Sanchez, A., O. Cast, and M.L. Miller, *Comprehensive Benchmarking and Integration of Tumor Microenvironment Cell Estimation Methods.* Cancer Res, 2019. **79**(24): p. 6238-6246.
12. Yoshihara, K., et al., *Inferring tumour purity and stromal and immune cell admixture from expression data.* Nat Commun, 2013. **4**: p. 2612.
13. Consortium, G.T., *The Genotype-Tissue Expression (GTEx) project.* Nat Genet, 2013. **45**(6): p. 580-5.
14. Levine, K.M., et al., *FGFR4 overexpression and hotspot mutations in metastatic ER+ breast cancer are enriched in the lobular subtype.* NPJ Breast Cancer, 2019. **5**: p. 19.
15. Zhu, L., et al., *Metastatic breast cancers have reduced immune cell recruitment but harbor increased macrophages relative to their matched primary tumors.* J Immunother Cancer, 2019. **7**(1): p. 265.
16. J, B., *seqsetvis: Set Based Visualizations for Next-Gen Sequencing Data*. 2021.
17. Love, M.I., W. Huber, and S. Anders, *Moderated estimation of fold change and dispersion for RNA-seq data with DESeq2.* Genome Biol, 2014. **15**(12): p. 550.
18. Federico, A. and S. Monti, *hypeR: an R package for geneset enrichment workflows.* Bioinformatics, 2020. **36**(4): p. 1307-1308.
19. Subramanian, A., et al., *Gene set enrichment analysis: a knowledge-based approach for interpreting genome-wide expression profiles.* Proc Natl Acad Sci U S A, 2005. **102**(43): p. 15545-50.
20. Zeng, Q., et al., *Synaptic proximity enables NMDAR signalling to promote brain metastasis.* Nature, 2019. **573**(7775): p. 526-531.

21. Hanzelmann, S., R. Castelo, and J. Guinney, *GSVA: gene set variation analysis for microarray and RNA-seq data.* BMC Bioinformatics, 2013. **14**: p. 7.
22. Shah, O.S., et al., *Identifying Genomic Alterations in Patients With Stage IV Breast Cancer Using MammaSeq: An International Collaborative Study.* Clin Breast Cancer, 2021. **21**(3): p. 210-217.
23. Smith, N.G., et al., *Targeted mutation detection in breast cancer using MammaSeq.* Breast Cancer Res, 2019. **21**(1): p. 22.
24. Merino, G.A., et al., *TarSeqQC: Quality control on targeted sequencing experiments in R.* Hum Mutat, 2017. **38**(5): p. 494-502.
25. Masica, D.L., et al., *CRAVAT 4: Cancer-Related Analysis of Variants Toolkit.* Cancer Res, 2017. **77**(21): p. e35-e38.
26. Chakravarty, D., et al., *OncoKB: A Precision Oncology Knowledge Base.* JCO Precis Oncol, 2017. **2017**.
27. Huang, L., et al., *The cancer precision medicine knowledge base for structured clinical-grade mutations and interpretations.* J Am Med Inform Assoc, 2017. **24**(3): p. 513-519.
28. Thorvaldsdottir, H., J.T. Robinson, and J.P. Mesirov, *Integrative Genomics Viewer (IGV): high-performance genomics data visualization and exploration.* Brief Bioinform, 2013. **14**(2): p. 178-92.
29. Basudan, A., et al., *Frequent ESR1 and CDK Pathway Copy-Number Alterations in Metastatic Breast Cancer.* Mol Cancer Res, 2019. **17**(2): p. 457-468.
30. Jambal, P., et al., *Estrogen switches pure mucinous breast cancer to invasive lobular carcinoma with mucinous features.* Breast Cancer Res Treat, 2013. **137**(2): p. 431-48.
31. Sikora, M.J., et al., *Invasive lobular carcinoma cell lines are characterized by unique estrogen-mediated gene expression patterns and altered tamoxifen response.* Cancer Res, 2014. **74**(5): p. 1463-74.
32. Qin, Y., et al., *Inhibition of histone lysine-specific demethylase 1 elicits breast tumor immunity and enhances antitumor efficacy of immune checkpoint blockade.* Oncogene, 2019. **38**(3): p. 390-405.
33. Liang, C.C., A.Y. Park, and J.L. Guan, *In vitro scratch assay: a convenient and inexpensive method for analysis of cell migration in vitro.* Nat Protoc, 2007. **2**(2): p. 329-33.
34. Tasdemir, N., et al., *Comprehensive Phenotypic Characterization of Human Invasive Lobular Carcinoma Cell Lines in 2D and 3D Cultures.* Cancer Res, 2018. **78**(21): p. 6209-6222.
